# BRAin-wide Neuron quantification Toolkit reveals strong sexual dimorphism in the evolution of fear memory

**DOI:** 10.1101/2022.09.12.507637

**Authors:** Alessandra Franceschini, Giacomo Mazzamuto, Curzio Checcucci, Lorenzo Chicchi, Duccio Fanelli, Irene Costantini, Maria Beatrice Passani, Bianca Ambrogina Silva, Francesco Saverio Pavone, Ludovico Silvestri

## Abstract

Fear responses are functionally adaptive behaviors that are strengthened as memories. Indeed, a detailed knowledge of the neural circuitry modulating fear and fear memory could be the turning point for the comprehension of this emotion and its pathological states.

A comprehensive understading of the neural circuits mediating memory encoding, consolidation, and retrieval over time presents the fundamental technological challenge of analyzing activity in the entire brain with single-neuron resolution. In this context, we developed the BRAin-wide Neuron quantification Toolkit (BRANT) for mapping whole-brain neuronal activation at micron-scale resolution, combining tissue clearing, high-resolution light-sheet microscopy, and automated 3D image analysis. The robustness and scalability of this method allowed us to quantify the evolution of neuronal activation patterns across multiple phases of memory in mice. This approach highlighted a strong sexual dimorphism in the circuits recruited during memory recall, which had no counterpart in the behaviour. The methodology presented here paves the way for a comprehensive functional characterization of the evolution of fear memory.

## INTRODUCTION

Fear responses are functionally adaptive behaviors that can be induced by direct encounter with a threat or with situations previously associated with a threat. Indeed, given its high survival value, the capability to remember potential threats is highly conserved across species (Kindt 2014). Although ‘fear’ refers to a human emotion, with a specific connotation in our minds not directly accessible in other animals, we can nevertheless study defensive responses induced by aversive events in animal models using standard behavioral paradigms like contextual fear conditioning (CFC) and inhibitory avoidance (IA) (Izquierdo, Furini, and Myskiw 2016). Fear induces many changes at different levels, from molecular and cellular to circuit ones (Josselyn, Köhler, and Frankland 2015; Josselyn and Frankland 2018). These permanent changes represent the physical trace of memory, commonly referred to as ‘engram’. In the last years, neuroscientists have made many steps forward in the knowledge of the molecular and cellular mechanisms underlying memory formation, consolidation, and retrieval (Johansen et al. 2011; Tonegawa, Morrissey, and Kitamura 2018; Josselyn and Frankland 2018; Orsini and Maren 2012). Furthermore, several brain areas, most prominently the hippocampus, the amygdala, and the prefrontal cortex, have been identified as important centers for memory processing (Izquierdo, Furini, and Myskiw 2016). However, several studies highlighted the involvement of many other regions (Pezzone et al. 1992; Moaddab, Ray, and McDannald 2021; Silva et al. 2021), supporting the hypothesis that memory is distributed and dispersed across the entire brain; in this perspective, the study of a single or few brain areas limits the comprehension of mechanisms underlying fear memory. The study of limited brain regions leads to a fragmented view that makes the global vision rarely interpretable; therefore, many significant questions about fear mechanisms and dynamics have yet to be explained.

The paucity of studies addressing fear memory substrates at the brain-wide level is mainly due to the technical limitations in large-scale analysis of neuronal activity. From a methodological point of view, understanding how neuronal networks drive this type of memory requires techniques for whole-brain activation mapping. In this respect, a promising strategy is to tag activated neurons *in-vivo* through immediately early gene (IEG) approaches and image them subsequently *ex-vivo* by 3D optical tools such as light-sheet microscopy (LSM). The coupling of clearing techniques, like CLARITY (Chung et al. 2013), iDISCO (Renier et al. 2014), or CUBIC (Susaki et al. 2014), with LSM has enabled the brain-wide mapping of cFos positive neurons activated in different behavioral contexts (Renier 2016, Roy 2022, Susaki 2014). However, IEG transgenic tools (Guenthner et al. 2013; DeNardo et al. 2019; Denny et al. 2014; Franceschini et al. 2020) label the entire neuronal cell, including axons and dendrites, making image analysis more challenging as compared to anti-cFos immunostaining which is confined to the nucleus (Franceschini et al. 2020). In this respect, sub-cellular resolution is needed to disentangle the contribution from neuronal processes and somata. Since the data size scales with the third power of the sample linear dimensions, even a moderate increase in spatial resolution (e.g., 2x) leads to a large increase of in the amount of data to be processed (8x in our example). Unfortunately, available analysis tools (Renier et al. 2016) are not capable of routinely handling terabyte-sized datasets, limiting brain-wide activation analysis to anti-cFos immunostaining. On the other hand, the ability to quantify cells labeled through IEG-based transgenic strategies would enable further investigation on the same animal after tagging of activated neurons (Roy et al. 2022; DeNardo et al. 2019).

From a clinical point of view, the study of fear memory requires scalable methods to analyze large cohorts of subjects. This is necessary to clarify how different neuronal circuits are disrupted in disease states such as in post-traumatic stress disorder (PTSD). Indeed, understanding how memory works and changes over time would imply analysis of multiple time points and of diverse experimental classes.

The lifetime prevalence of PTSD is about 10–12% in women and 5–6% in men (Charak et al. 2014). Women have a two/three times higher risk of developing PTSD compared to men (Christiansen and Hansen 2015). In spite of this striking difference in prevalence and risk between male and female subjects, sex difference studies are only partially considered in the field of neuroscience, since researchers typically favor male mice rather than female mice in their studies, with a ratio of 5.5 to 1 (Zucker and Beery 2010). This disproportion does not consider the sex statistical differences in the lifetime of many pathologies, suggesting serious implications for healthcare in women. The factors that influence the different lifetimes in PTSD vary from biological to psychological ones (Christiansen and Hansen 2015). Also in the context of whole-brain mapping, the few studies published hitherto focused only on the male population (Cho, Rendall, and Gray 2017; Roy et al. 2022; Vetere et al. 2017; Wheeler et al. 2013; Bonapersona et al. 2022).

Here, we present BRANT (BRAin-wide Neuron quantification Toolkit), a new pipeline for whole-brain mapping, exploiting TRAP mice (Guenthner et al. 2013), high-resolution LSM, and terabyte-scale image processing. Using BRANT, we analyze the evolution of whole-brain neural circuits recruited upon aversive memory in females and males. We find strong sexual dimorphism in the evolution of whole-brain networks underlying fear memory, both at the level of activation patterns and of functional connectivity.

## RESULTS

### 1. BRANT enables scalable, high-resolution analysis of neuronal activity patterns in behaviorally-relevant cohorts

Any method aiming at complementing behavioral analysis with physiological or anatomical data must be applicable to dozens of samples, allowing its use in a statistically significant number of animals for each behavioral group. With this important constraint in mind, we developed BRANT, a scalable and user-friendly pipeline for 3D analysis of whole-brain activation patterns, that combines FosTRAP transgenic strategy (Guenthner et al. 2013), CLARITY/TDE (Costantini et al. 2015), high-resolution RAPID-enabled LSM (Silvestri et al. 2021) and 3D automated data analysis (Frasconi et al. 2014) (**Fig.1A**). BRANT was validated using a classical paradigm, i.e. step-through passive IA. After the behavioral task, mice are injected with 4-hydroxytamoxifen (4-OHT) to drive Cre-mediated recombination in cFos-expressing neurons (Guenthner et al. 2013).

**Figure 1:**
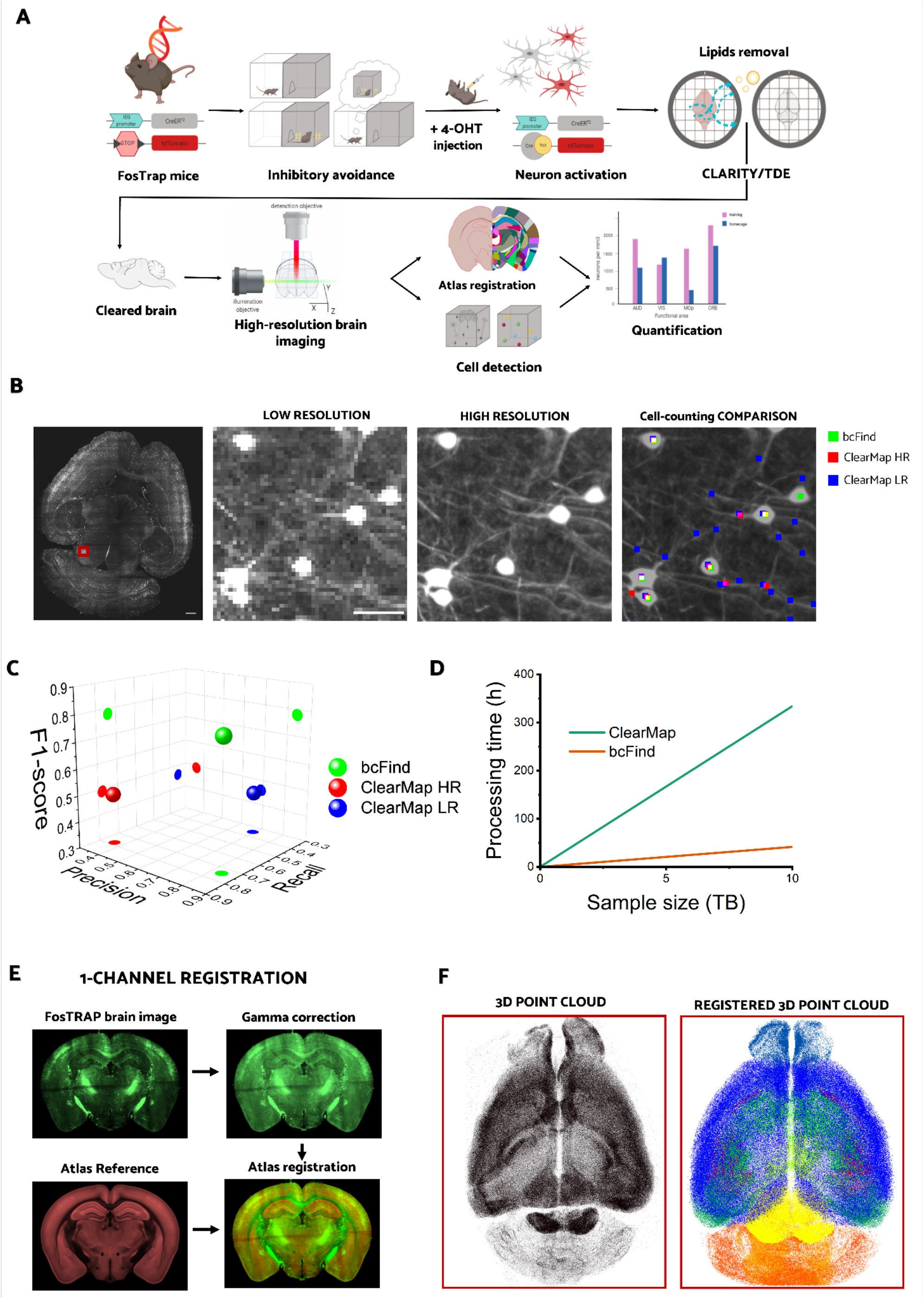
Experimental pipeline. (A) Schematic representation of various steps constituting the experimental pipeline. FosTrap mice underwent the step-through IA task. Injection of 4-OHT drives permanent expression of tdTomato in activated neurons. After perfusion, all brains were processed with CLARITY/TDE and imaged with a high-resolution RAPID-enabled LSM. High-resolution imaging was fundamental for cell detection and 3D automated analysis. All brains were registered to the Allen Brain Atlas. tdTomato-positive neurons are automatically detected and quantified across all behavioral groups. (B) Virtual slice extracted from a whole-brain tomography (left). The red square identify a small region zoomed in the other subpanels. Low- and high-resolution zoom-ins, corresponding to a voxel size of 4×4×4 µm^3^ and 0.65×0.65×2 µm^3^, respectively (center). Results of cell counting in this subvolume obtained with bcFind and ClearMap (right). (C) Comparison of the localization performance (precision, recall and F-1 score) of bcFind and ClearMap at low resolution (ClearMap LR) and high resolution (ClearMap HR). (D) Data throughput of bcFind and ClearMap in 3D automated analysis. (E) One channel image registration to the Atlas. Gamma correction is applied to the FosTrap image. Image with high contrast is used for the Atlas registration. (F) The point cloud obtained with bcFind (left) is finally warped to the reference atlas (right). In the right image, points are colored according to the brain region they belong to. Scale bar, 50 µm.

Importantly, since the reporter fluorescent protein (tdTomato) is not confined to the soma but is expressed in the entire neuron, high-resolution imaging is required to disentangle dense environments (**Fig.1B**). To this aim, 3D whole brain reconstructions were achieved by optimizing clearing and imaging protocols. All brains were made optically transparent using a modified CLARITY/TDE protocol allowing moderate expansion of tissue (27% linearly) (Di Giovanna et al. 2019). This isotropic change in sample size offered the possibility to discriminate single cells in densely labeled structures. Samples were then imaged using a custom-made LSM with confocal line detection, able to reconstruct mesoscopic samples with microscale resolution (Müllenbroich et al. 2015). Through a system for real-time stabilization of light-sheet alignment, called RAPID, we were able to maintain high resolution across the entire sample volume (Silvestri et al. 2021). Since RAPID operates simultaneously with image detection, no acquisition overhead is introduced.

Each brain reconstruction comprises about 16 TeraBytes of raw data with a voxel size of 0.65×0.65×2 μm^3^. This data size is incompatible with a cohort study as it would require storage capabilities in the order of 1 PetaByte for a single study. For this reason, the acquired datasets were first compressed by a factor 20 using the 16-bit lossy JPEG-2000 format, thus reducing disk usage while still retaining overall good image quality and detail level (**Video V1**). Images were then stitched using ZetaStitcher (https://lens-biophotonics.github.io/ZetaStitcher/), a custom-made Python software for large volumetric stitching specifically developed for light-sheet microscopy. An important feature of ZetaStitcher is VirtualFusedVolume, an application programming interface (API) that provides seamless and effective access to high-resolution data by simply providing the spatial coordinates of the sub-volume of interest within the virtually fused volume. In this way, large volumes can be programmatically processed in smaller chunks in a distributed environment and without user intervention, a key requirement to process the large datasets produced by high-resolution LSM. Fluorescently labeled activated neurons were detected using BrainCell Finder (bcFind) (Frasconi et al. 2014). This 3D automated analysis relies on a deep learning approach, based on a U-net architecture, widely used in the image-analysis field (Ronneberger, Fischer, and Brox 2015), to recognize specific structures in complex data sets. Indeed, since tdTomato is expressed also in neuronal processes, the application of standard methods based on thresholds and blob detectors is not feasible (**Fig. 1B**). The task of our U-net is to transform raw images, containing many disturbing objects (like axons, dendrites, and vessels), into ideal images that include only small spheres at the location of neuronal bodies. In the images transformed by the U-net, a basic blob detector algorithm can then effectively localize the fluorescent cell bodies. The performance of this method was evaluated on 3’383 manually labeled cells, obtaining precision = TP/(TP+FP) equal to 0.84, recall = TP/(TP+FN) 0.74 and F1-score = 0.78. F1 is defined as the harmonic mean of precision and recall, 1/F1 = (1/P + 1/R)/2. We compared the accuracy of bcFind with that obtained using ClearMap, a widely used software for cFos+ cell detection in cleared brains (Renier et al. 2016). To simulate the typical application settings of the method, we tested it also on images downscaled to a standard voxel size of 4×4×4 µm^3^ (**Fig. 1C**). ClearMap analysis showed poor performance both with high- (precision 0.36, recall 0.78, F1 0.49) and low-resolution data (precision 0.69, recall 0.35, F1 0.46). This is not surprising since the method has been developed to detect spherical-like objects (like cFos-stained nuclei) and not complex neuronal cells.

Given that raw data amount to about 16 TeraBytes per brain, another fundamental feature is computational scalability. bcFind can be operated in batch mode on a computing cluster, enabling image processing at a rate which is ultimately limited only by computational power. In our case, using a cluster with 8 GPUs (NVIDIA GeForce RTX 2080Ti) we were able to process data at about 240 GB/h, about 1 order of magnitude faster than the reported speed of ClearMap (Renier et al. 2016) (**Fig. 1D**). Importantly, this is not a comparison about the absolute speed of both algorithms – i.e. these data does not mean that bcFind is faster than ClearMap when using the same resources – but a comparison about the capabilities of the two software to effectively run on large-scale parallelized environments – meaning that in practical settings bcFind is faster when used on TB-sized datasets.

To compare cell counts across multiple subjects, 3D brain datasets are spatially aligned to the Allen Brain Reference Atlas using Advanced Normalization Tools (ANTs) (Avants et al. 2011). Differently from previous reports, we did not acquire a secondary channel for atlas registration but applied a strong gamma correction to reduce the contrast between tissue autofluorescence and the signal from fluorescent protein (**Fig. 1E-1F**). We found a registration accuracy of about 300 µm (see Methods), which is comparable to that obtained by ClearMap (Renier et al. 2016). Notably, the approach implemented here reduce the cost and complexity of the microscope used, since only one camera is needed in the microscope and the size of acquired datasets is not duplicated. Of note, all brain areas were included and grouped for this study, manually selecting 45 macro areas. We decided to use larger areas (the Allen Reference Atlas has more than 1’000 subregions) based on the measured registration accuracy, to avoid misassignment of cells.

In conclusion, the combination of fast high-resolution imaging and scalable 3D analysis for processing sub-cellular information is the core of this pipeline, enabling quantification of active neuronal ensembles in behaviorally-relevant cohorts (**Videos V2-V3**).

### 2. BRANT analysis reveals sexually dimorphic patterns of brain activation in different phases of aversive memory

BRANT was applied to study the time evolution of neuronal activity over the course of fear memory **(Fig. 2A)**. In order to answer our biological question, we used step-through IA as a behavioral paradigm in which mice learn to associate a particular context (i.e. white/black box) with an aversive event (i.e. a mild foot shock). Differently from the standard CFC, IA implies decision-making since mice can decide whether to step into the dark compartment when they are subsequently tested for memory retention (Izquierdo, Furini, and Myskiw 2016). Indeed, the latency time to enter the dark compartment is used as a direct measurement of memory. This task relies on a single training session and produces robust memory that is easily quantifiable and long-lasting. By using IA, we selected three experimental groups based on three different memory phases describing the evolution of fear memory, from encoding to retrieval: *training group* was selected to study fear encoding while test *groups* at *24 hours* (24h) and *7 days* (7d) after training were selected to explore the recent and long-term fear memory retrieval, respectively **(Fig. 2A)**. As expected, latency times of training groups (in which the latency to enter the shock compartment is measured in the habituation period) were significantly different compared with the latency times of the respective test groups. Statistical analysis indicate that latency times were influenced only by the experimental class and not by sex (2-way ANOVA, followed by Bonferroni’s post-hoc comparisons tests) **(Fig. 2B)**. Indeed, there was no significant difference in training performances in any group examined or between various test groups **(Fig. 2B)**. This indicates that all mice, independently from their sex, formed a memory of the training experience even though the shock intensity (0.3 mA) was weaker than the classically used (Canto-de-Souza and Mattioli 2016; Benetti et al. 2015). The value of shock intensity was selected within a range in which behavioral outcome at retrieval shows distinct inter-subject variability, with the aim to point out the contribution of neuronal activation resulting from the decision-making process. (**Fig. S1**). Although the results from IA did not highlight any differences between male and female groups, the quantification of activated neurons (cFos) in 45 brain regions (for a complete listing of brain regions, see **Table S1**) revealed a strong sexual dimorphism in the underlying neuronal activity pattern. To obtain a statistically robust comparison of the activation patterns between male and female subjects, we performed mean-centered partial least squares analysis (PLS) (Krishnan et al. 2011). This method identifies a set of latent variables (‘contrasts’) in the space formed by the different experimental conditions, and a corresponding set of ‘saliences’, i.e. the contribution of different brain areas in differentiating the samples between the different contrasts. Notably, differently from pairwise comparison of cell counts between different experimental classes, this method does not require correction for multiple sample comparisons, resulting in overall higher statistical power (Krishnan et al. 2011).

**Figure 2:**
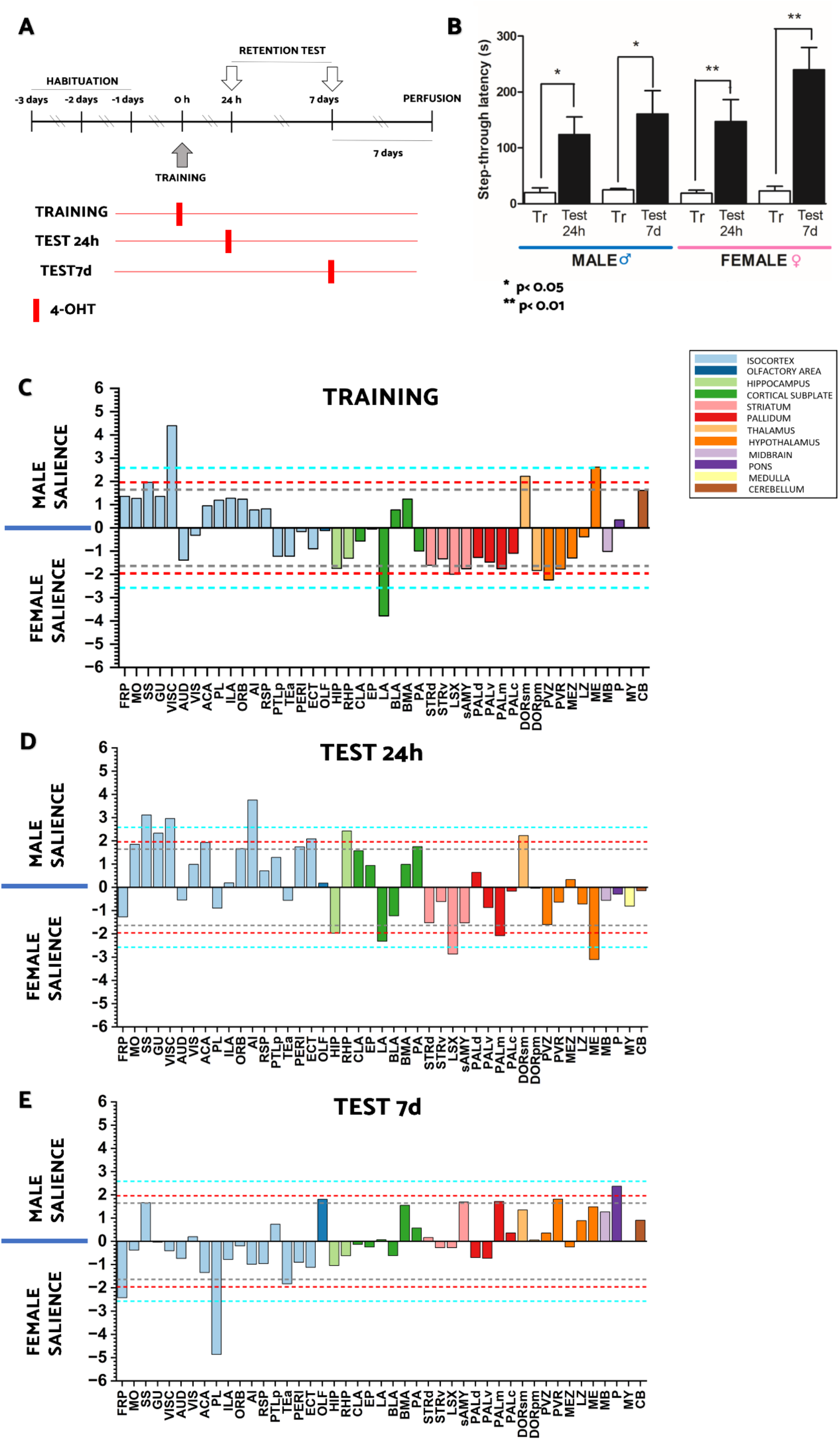
Behavioral task and neuronal activity analysis. (A) Schematic representation of the experimental paradigm. Animals are handled for three days before training. During the training session, all mice receive a foot-shock (0.3 mA, shock duration 2 s, delay after closing door 0.5 s). Memory retention test is performed 24h or 7 days after training. By the 4-OHT injection at three time points, mice were divided in three experimental classes: training; test 24h, test 7d (B) The comparisons of acquisition and retention latency times are analyzed by 2-way ANOVA, followed by Bonferroni’s post-hoc comparisons tests (* p < 0.05, ** p < 0.01, training n=6 males and n=4 females, test 24h n=5 males and n=6 females and test 7d n=4 males and n=4 females). Data are expressed as means ± SEM of 4 to 6 animals for each group. (C-E) PLS analysis of cFos expression between the two sexes in all experimental classes: training (C), test 24h (D), test 7d (E). Salience scores, normalized by standard deviation calculated with bootstrap (right), identify regions that maximally differentiate between these conditions. The red, gray and blue lines reflect respectively a salience score of 1.64 (p < 0.1), 1.96 (p < 0.05) and 2.58 (p < 0.01).

The vectors of PLS contrasts that discriminate between sexes have different structure at different time points (training, test 24h and test 7d after training), revealing distinct patterns of cFos expression between males and females in our experimental groups (**Fig. 2C-2E**).

Using this unbiased analysis, a strong sexual dimorphism in the recruitment of several brain areas in each memory phase was observed. This dimorphism emerges not only in regions classically considered to be involved in aversive memory such as prefrontal cortex, amygdala and hippocampus, but also in brain areas less known for their contribution in this function such as pallidum, striatum, thalamus and other subcortical regions. The differences of cFos expression (**Fig. 2C-2E**) in all brain regions seem to fade over time, from training group to test 7d group. Indeed, 7 days after training, there is an alignment of cFos patterns between sexes with only 3 brain areas (FRP, PL, P) being significantly different with p<0.05 (**Fig. 2E**). At training and test 24 hours after training, male mice showed an increased cFos expression in the cerebral cortex (SS, GU, VISC, AI, ECT) while female mice in subcortical regions, with the exception of the thalamus that is always more activated in male group at all time points (**Fig. 2C-2D**). In detail, at 7 days after training, male mice are associated with an increased cFos expression in the pons and other evolutionary older brain regions while female mice are associated with an increased cFos expression in the associative cortex (FRP and PL) (**Fig. 2E**). An interesting case is represented by the median eminence that is more activated at training in males while at test 24h in females (**Fig. 2C-2D**).

These data indicate that males and females activate, on average, different neuronal patterns in distinct phases of aversive memory, highlighting a strong sexual dimorphism which is, notably, not reflected in behavioral differences.

### 3. Neuronal activity correlates with behavioral outcome in different brain regions in females and males

BRANT application showed how activity patterns involved in fear memory differ in male and female mice. The results obtained by PLS, based on tdTomato^+^ neuron quantifications in each brain region, showed the areas which account more for the difference in activity between experimental groups, each one corresponding to a specific memory phase. Thus, this analysis highlights variations between different experimental classes at population level, but is not appropriate to investigate the inter-subject variability inside the same experimental group. Indeed, latency times were significantly dispersed, underlining variable behavioral responses between different mice (**Fig. 2A, Fig. S1**). To investigate whether neuronal activation of specific areas can be accounted for the behavioral variability observed, we performed a cross-correlation analysis between regional counts and latency times (**Fig.3**). This analysis led to the identification of strong correlations or anti correlations (p<0.01) for different areas in different experimental classes. Importantly, brain regions that correlate with behaviour are different for both sexes, underlining a sexual dimorphism also in this type of analysis (**Fig.3**). As expected, amygdala, hippocampus and prefrontal cortex are correlated with the step-through latency times, but also the activation of areas such as pallidum, striatum, pons, and other regions less known to be involved in fear memory, correlate with the times mice spent in the bright cage before stepping through in the dark compartment.

**Figure 3:**
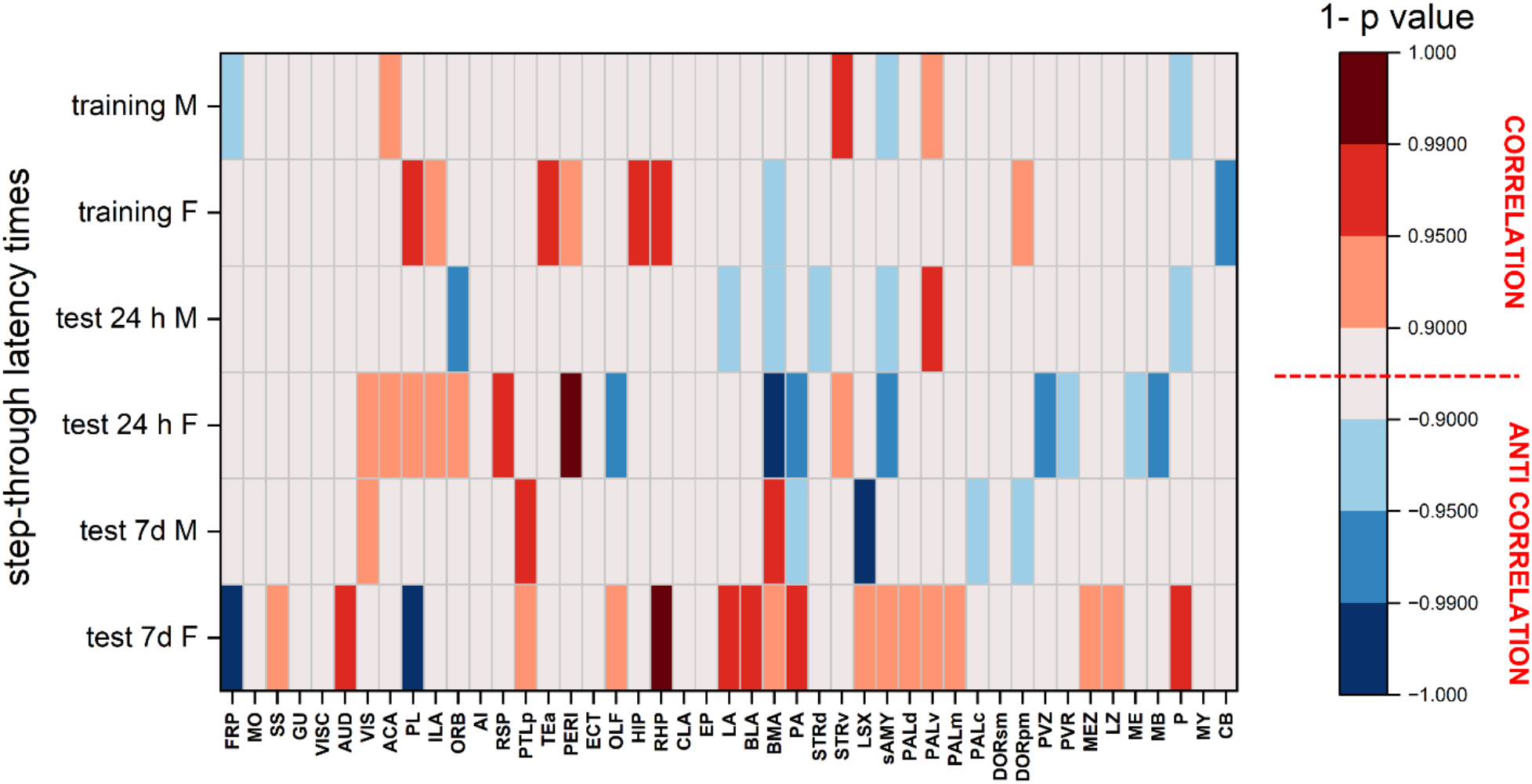
Pair-wise correlation of latency time with neuronal activity in each brain area. Pearsons correlation heat maps. Here heat maps represent the correlation between the number of tdTomato^+^ neurons in each region and the latency time mice spend to step-through into the dark compartment. On the right side of the plot, different colors correspond to different significant degrees (1 - p values).

For male and female training groups, latency times are not directly related to fear but instead to the exploration of the environment. For this reason, in line with the literature, the brain areas more correlated with these latency times are: TEa, HIP and RHP, relevant for specific contextual information (Bucci and Robinson 2014; Izquierdo, Furini, and Myskiw 2016; D. M. Smith and Bulkin 2014), STRv which supports rapid discrimination of uncertain threat that is necessary for the first contextual exploration (Ray et al. 2020), PL that regulate fear expression (Giustino and Maren 2015) and CB that participates in movement and it is an important autonomic control center as part of an integrated network regulating fear (Sacchetti et al. 2002). Notably, CB is anticorrelated with training latency times, suggesting that its activation leads to reduced explorative behavior. Here, male subjects show a significant correlation between behavior and neuronal activation only in STRv. On the other hand, in female subjects strong correlation is observed in HIP, RHP, PL, TEa, while a clear anticorrelation is found in CB.

Latency times during test sessions are usually considered in IA a measure of fear memory retention (Izquierdo, Furini, and Myskiw 2016). For male mice, times measured at 24h show a strong correlation with PALv and an anticorrelation with ORB. This finding is in contrast with a recent study showing that neuronal firing in ventral pallidum is inhibited in presence of a potential danger (Moaddab, Ray, and McDannald 2021). However, that experiment was carried on in a different species (rat) and involved a different behavioral paradigm (cued fear conditioning). Nonetheless, these two results underline the important yet hitherto neglected role of PALv in fear memory. Female mice tested 24 hours after training showed strong correlation with RSP and PERI activity. Previous studies highlighted the crucial role played by RSP in the formation of the contextual fear memory (Kwapis et al. 2015), consistently with our findings that the more this region is activated the longer is the IA latency time. OLF, PA which is areas belonging to the amygdala, well known for its role in fear memory (Fareri and Tottenham 2016; Hoffman et al. 2014; Sacchetti et al. 2002; Kitanishi and Matsuo 2017), PVZ that is an important autonomic control center in response of stress exposure (Herman and Tasker 2016), sAMY which supports affective evaluation and learning (Fareri and Tottenham 2016), and MB are instead characterized by a distinct anti-correlation with step-through latency.

In males, latency times 7 days after training show strong correlation with PTLp and anti-correlation with LSX. This finding confirms the involvement of LSX in memory and emotional responses: indeed previous reports show that this region is activated during aversive situations (Pezzone et al. 1992; Beck and Fibiger 1995; Duncan, Johnson, and Breese 1993) and projects to brain regions involved in the behavioral and cardiovascular responses to aversive stimuli (LeDoux et al. 1988). Female subjects show a completely different pattern of correlation with behavior. Latency times are significantly anti-correlated with activation of FRP and PL, areas that are significant sexually dismorphic in the PLS analysis too. Notably, a recent study highlighted the role of FRP in decision making as that of the whole prefrontal region (Lv et al. 2021). While the correlation of latency times are with RHP, AUD, P and all amygdalar regions, areas that are well known to be involved in fear (Fareri and Tottenham 2016; Hoffman et al. 2014; Sacchetti et al. 2002; Kitanishi and Matsuo 2017). Interestingly, at 7d retrieval, PA and LSX shows an opposite correlation between sexes. On the other hand, BMA is the only exception to a general sexual dimorphism which, for both sexes, anti-correlate 24 hours after training and correlate at 7 days after. Generally, the behavioral relevance of BMA at retrieval, highlighted by Pearson correlation analysis between regional activation and behavioral responses, is stronger for females compared to males.

### 4. Network analysis shows strong sexual dimorphism in the evolution of functional connectivity

Widely accepted theories affirm that memory is distributed across multiple brain regions that are functionally connected (Roy et al. 2022; Frankland and Bontempi 2005). To estimate average functional connectivity in different behavioral groups, we performed a Pearson cross-correlation analysis between normalized activation counts of different brain areas, among subjects of the same experimental group (**Fig. S2**). Considering only the correlations and anticorrelations with p<0.05 (**Fig. S3**), we identified a network of brain areas that shows a statistically significant tendency to be active together (for correlations) or in a mutually exclusive manner (for anticorrelations) (**Fig. 4**). Thus, brain areas are the nodes of this network, whilst the edges represent putative functional connections identified by super-threshold cross-correlation.

**Figure 4:**
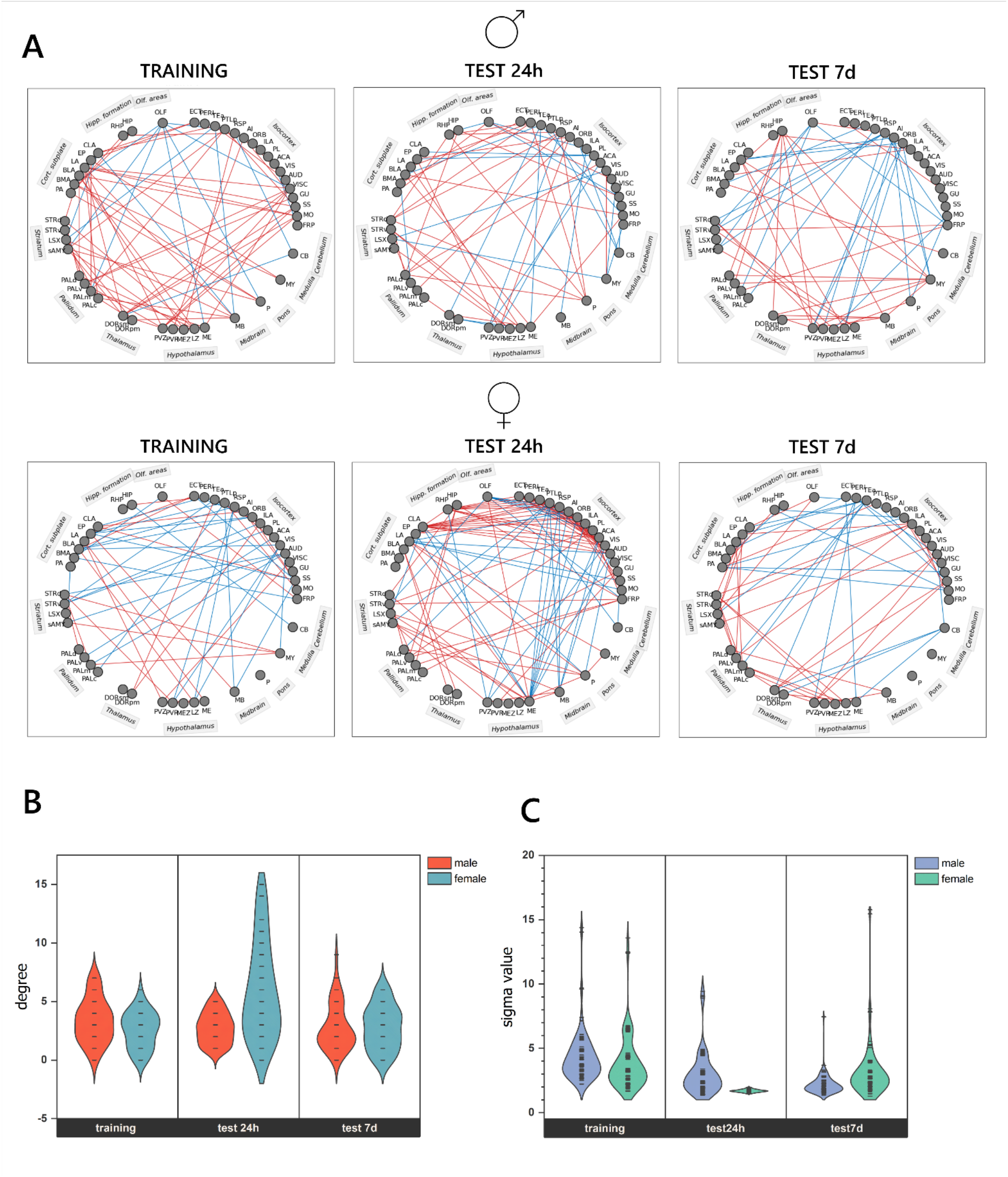
Fear memory networks in male and female mice. (A) Network graphs were generated by considering only the strongest correlations (see Methods). Grey circles represent the 45 selected regions **(Listed in Table S1**). The lines between the nodes represent significant positive (red) or negative (blue) correlations (see methods for thresholding details). (B) Violin plot of the connectivity of all the nodes in the networks shown in (A). Violin plots of sigma values for the networks shown in (A), calculated for N=100 generations of random graphs (see methods).

Interestingly, females and males activity networks showed a different evolution over time (**Fig. 4A**). In order to quantitatively analyze differences in organization, we compared a number of connectivity features in male and female networks. The distribution of nodes degree (Hallquist and Hillary 2019) was similar for the two sexes at both training and test 7d, while 24 hours after training the female network was characterized by a large increase in connectivity (**Fig. 4B**). Interestingly, in male mice we observe a decrease of the relative contribution of positive correlations from training (82%) to test 7d (64%) with a minimum at test 24h (59%), while females show the opposite trend (50% at training, 63% at test 7d) with a peak at 24h after training (74%, **Fig. S4**). If we restrict our analysis to the isocortex, we observe that the intraregional connectivity (normalized on the global average node degree) follows a trend similar to the global connectivity. These results suggests that engram migration to the cortex, which is a cornerstone of system consolidation theories (Albo and Gräff 2018; Frankland and Bontempi 2005; Wheeler et al. 2013), occurs with different spatio-temporal patterns in males and females, but both sexes tend to converge to the same network structure (**Fig. S5**).

The sexual dimorphism observed in network evolution is confirmed by small-world analysis. For each network, we evaluated the small world coefficient σ of the giant component (i.e., isolated nodes were not considered), defined as (C/Cr) / (L/Lr), C and L being the average clustering coefficient and the average shortest path lengths of the graph, respectively. Cr and Lr are the corresponding values for an equivalent random graph. σ is equal to one in graphs with a structure similar to random ones, whereas it is higher than one when the network is “small-world”, i.e. when non-neighbouring regions are separated by a small number of steps compared to a random network. At training and test 7d, we observed a network presenting small word features for both sexes. Conversely, the network of the *24h group* largely resembles a random graph for the female group but not for males (**Fig. 4C**). This finding supports the hypothesis that the circuitry of male and female mice evolves towards an organized cortical network following different pathways.

### 5. Hubs of functional connectivity change over time in a sexually dimorphic manner

In addition to global properties of the connectivity graph, we assessed the role of different nodes by measuring their degree and betweenness. The degree of a node is the number of connections it has with the other nodes in the network, whereas betweenness is defined as the probability of finding the node in the shortest path between any randomly chosen pair of nodes in the network. In other words, the degree is a direct measure of the influence of the node in the network (the higher the degree, the more the activity of the node correlates with that of other nodes), whilst the betweenness states the importance of the node for connectivity (i.e. if a node with high betweenness is removed, many paths between other nodes are cut). We considered as hubs those nodes which are simultaneously above the 80 percentile of degree and betweenness values (**Fig. 5**). We found that region involvement as hubs shows distinct time evolution in males and females (**Fig. 6, Table S2**).

**Figure 5:**
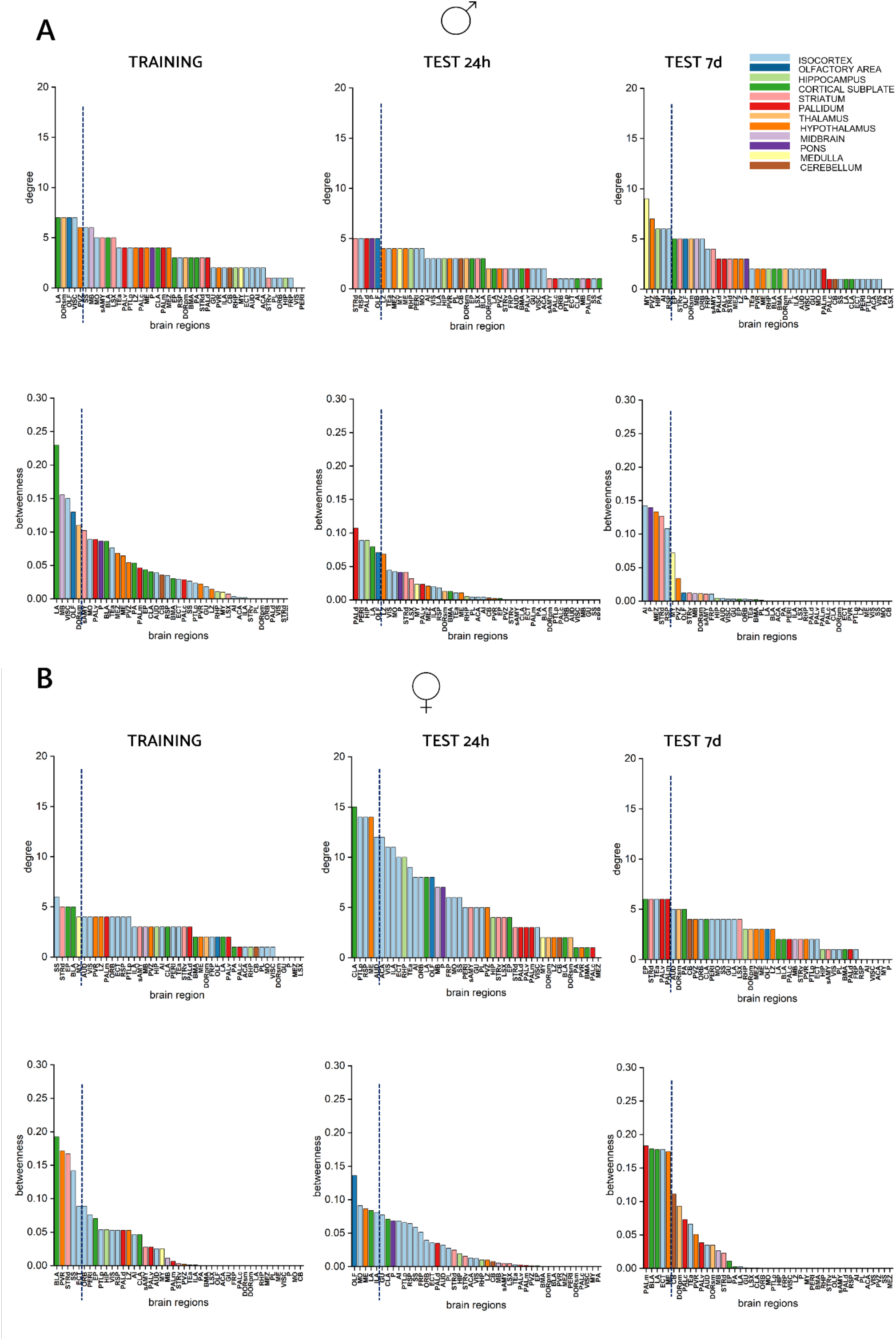
Identification of hubs in memory networks. Brain areas ordered according to degree or betweenness values in the three experimental time points for both male (A) and female (B) subjects. Vertical dashed lines denote the 90^th^ percentile. Nodes in the 90^th^ percentile for both degree and betweenness are considered as hubs.

**Figure 6:**
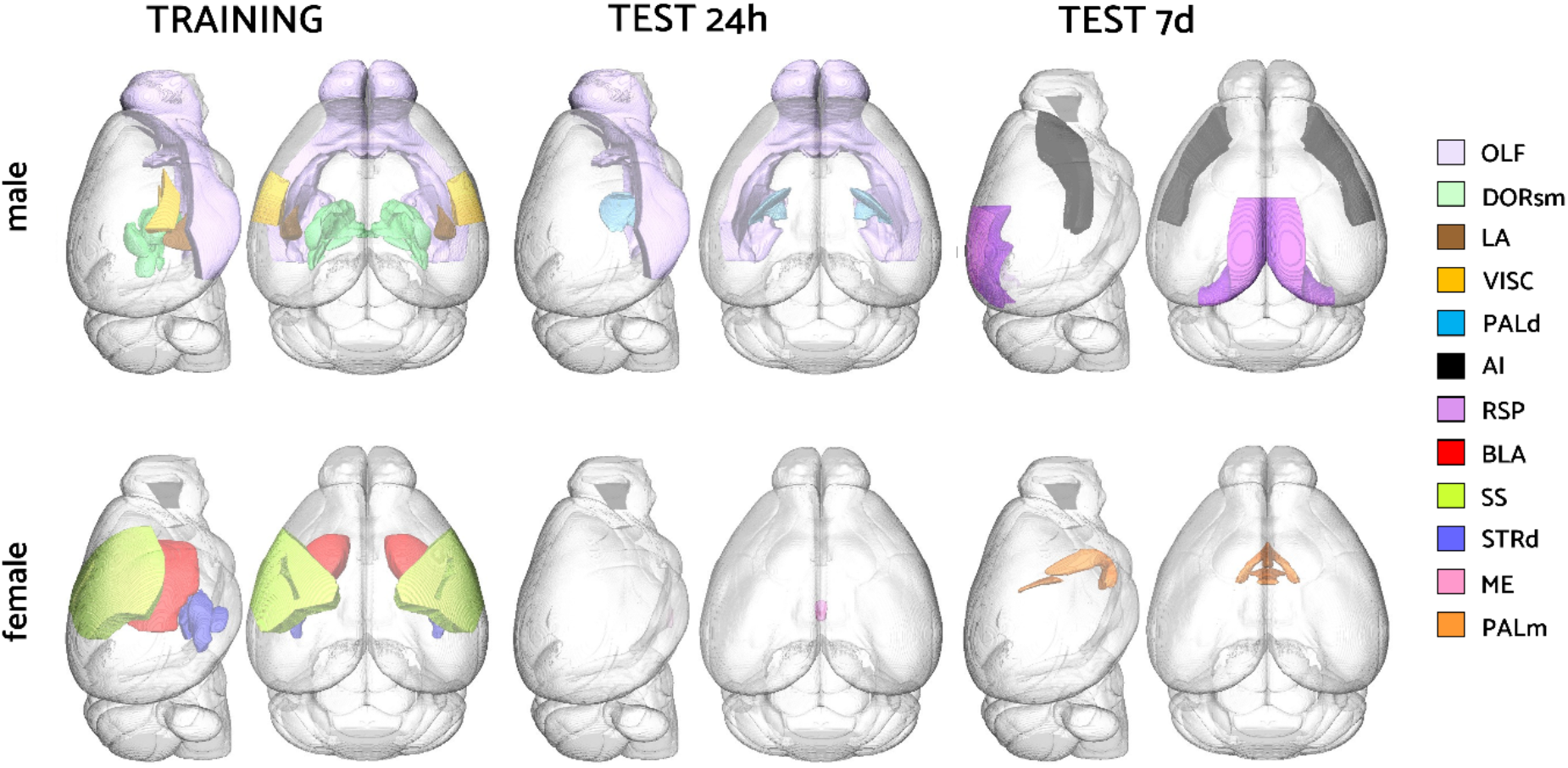
Spatio-temporal evolution of functional connectivity hubs. 3D rendering of a reference mouse brain where hubs of specific time points are depicted with different colors. Each color represents a single brain region according to **Table S1**.

In male subjects, we observed a hub evolution from subcortical areas at training (LA, DORsm and OLF) and test 24h (PALd and OLF) to associative corticices (RSP and AI) at 7 days after training. The visceral cortex, which is a hub at training, is a relevant exception to this general scheme. These results are in line with the standard theory of memory consolidation, where fear-processing neuronal circuits migrate from amygdala and thalamus to the cortex (Izquierdo, Furini, and Myskiw 2016). Conversely, in female subjects, network hubs persist in subcortical regions across the entire time span investigated, moving from BLA and STRd at training, ME at test 24h, and PALm at test 7d. Again, at training we found a cortical hub (SS) which, similar to what observed in male mice, is in a region processing proprioceptive stimuli, suggesting a role of these hubs in elaborating the unconditioned stimulus (foot shock).

These results are only partially in line with the PLS analysis. Indeed, most hubs are not significantly more active in males rather than in females, with the notable exception of VISC and ME, which are both hubs and preferentially activated in males at training and in females at test 24h, respectively. Notably, lateral amygdala is an hub for male subjects at training but is significantly more recruited in females at the same stage (Fig. 2C).

Consistently with this apparent uncoupling between and regional activation and functional connectivity role of brain areas, we observed a common feature of both male and female networks: hub regions for each memory phase are distinct from areas where activation significantly correlates with step-through latency times. Overall, these results suggest that hubs do not directly modulate behavior, but rather orchestrate downstream circuits mediating behavioral output.

## DISCUSSION

Memory, as many other cognitive processes, involves complex neuronal circuits that are distributed across the entire brain and change with time (Herry and Johansen 2014). Thus, from a technical perspective, it is important to have tools that allow scalable, unbiased, and quantitative analysis of brain activity at organ level yet with single-cell resolution. The classical procedure to reach this aim consists in serial sectioning of the tissue, immunostaining against cFos (or another IEG), and analysis of tissue slices (Kim et al. 2015). Although this protocol has been exploited to characterize the networks recruited during contextual (Wheeler et al. 2013) or tone fear conditioning (Cho, Rendall, and Gray 2017), a massive amount of work is required to process each murine brain, thus resulting impractical for routine use in many laboratories. Automated 3D imaging, such as serial two-photon sectioning (STP) (Ragan et al. 2012) and light-sheet microscopy (Mertz 2011), emerged in the last decade as a potential game changer for whole-brain analysis at cellular level, and indeed it has been exploited to map neuronal activation in different contexts (Renier et al. 2016; Kim et al. 2017), including fear memory (Roy et al. 2022; Bonapersona et al. 2022). However, STP is usually employed in a sampling scheme, failing to cover the entire tissue volume (Kim et al. 2017). On the other hand, LSM-based pipelines, like ClearMap, are typically limited to a resolution of several μm per voxel, preventing their use in densely labeled regions (**Fig. 1**). Here, we presented BRANT, a combination of high-resolution LSM and scalable computational analysis. On the optics side, the use of RAPID autofocusing guarantees high contrast and resolution across the entire sample (Silvestri et al. 2021). On the image processing side, the architecture provided by ZetaStitcher allows programmatic access to small chunks of data in an otherwise extra-large tissue volume (several TeraVoxels) for subsequent quantification, and the use of a deep learning method for cell detection (bcFind) enables superior accuracy when applied to neurons labeled in their entirety (compared to nuclear staining of anti-cFos immunohistochemistry). Overall, these innovations enable routinary and scalable analyses that were not possible using previously reported methodology. Importantly, BRANT is based on fully open-source software available on GitHub, and can thus be used and improved by the entire community in a collaborative spirit.

The lack of scalable methods for whole brain mapping has limited the analysis of brain-wide activation patterns in fear memory to a handful of studies (Cho, Rendall, and Gray 2017; Tonegawa, Morrissey, and Kitamura 2018; Bonapersona et al. 2022; Vetere et al. 2017; Wheeler et al. 2013). Most reports about fear memory refer to one or few nuclei or areas. Consequently, the obtained results are specific to those neuronal regions, which in turn makes it difficult to form an overview when integrating the data. Indeed, small changes in the experimental setting (from the behavioural task to the age and sex of animals) can lead to contrasting outcomes, making it difficult to obtain a comprehensive view of brain circuits underlying memory formation, consolidation, and retrieval. Additionally, this problem is even more accentuated considering the limited literature about female subjects and sex differences. Indeed, the majority of previous studies about sexual dimorphisms have mainly highlighted differences in brain anatomy (Kim et al. 2017; Bayless and Shah 2016; Simerly 2002; Cachero et al. 2010) and in cellular physiology (Florido et al. 2021). The identification of these anatomical or molecular differences is a useful starting point for the comprehension of neural circuits underlying sex-specific behaviors, but brain-scale functional studies are needed to disentangle the shape of different circuits recruited during the evolution of aversive memory.

In this context, our study demonstrated that fear memory is associated to the recruitment of sex-specific networks. It is noteworthy how this difference is emphasized during the 24h retrieval, suggesting that female mice augment their brain-wide functional connectivity state, to evolve quicker to a network similar to the starting conditions, with preferential involvement of different brain regions. Interestingly, the functional network of male mice changes over time, recruiting distinct regions but not increasing the number of nodes and connections at each memory phase. Overall, these data suggest that fear learning and retrieval are mediated by distinct subsystems in the two sexes challenging the use of a male-predominant literature for understanding aversive memory in females. Generally, this study could pave the way for a better understanding of those neuro-psychiatric disorders that exhibit sex differences in lifetime prevalence such as PTSD anxiety and depression (Bangasser and Valentino 2014; K. M. Smith and Dahodwala 2014; Werling and Geschwind 2013; Davies 2014; Li and Singh 2014).

Here, BRANT was applied to understand the time-evolution of brain circuits underpinning fear memory in an IA paradigm with a mild footshock. Due to the limited number of whole-brain studies about fear memory, and the lack of involvement of female subjects, comparing our results with previous reports (Wheeler et al. 2013; Cho, Rendall, and Gray 2017) is not straightforward because fear memory can be assessed in rodents by using diverse behavioral tasks. Indeed, the IA behavioral paradigm presented here might recruit a different circuitry with respect to context or auditory fear conditioning, since decision-making - to step through the gate towards the dark compartment - is an important component, absent in the above mentioned paradigms. In addition, using a mild foot shock (0.3 mA), we explored an aversive rather than traumatic memory. Nevertheless, our results confirm the unified engram complex theory (Roy et al. 2022), proving that memory evolution recruits dynamical networks across the entire brain.

In this fragmented context - both on the brain region studied and on the behavioral task used - the scalable and comprehensive analysis enabled by BRANT can play an important role. Indeed, this pipeline exploits well-described protocols and open-source software, offering to any lab the possibility to perform whole-brain activation analysis at high throughput, allowing cell-resolution brain mapping on behavioral cohorts. Here, this method highlighted a sexual dimorphism in the evolution of fear memory but it is readily adaptable for broader applications such as drug discovery or other biological purposes. The advantage of using micrometer resolution on a brain-wide scale and 3D automated analysis could revolutionize the way of studying brain connectivity and functions.

## METHODS

### ANIMALS

Male and female FosTRAP mice (B6.129(Cg)-Fos^tm1.1(cre/ERT2)Luo^/J × B6.Cg-Gt(ROSA)26Sor^tm9(CAG-tdTomato)Hze^/J) were used for this work (Guenthner et al. 2013). They were housed in groups of 3 or 4 with food and water ad libitum and were maintained in a room under controlled light and dark cycle (12/12 h; light starts at 7:00 AM), temperature (22 ± 2 °C), and humidity (55 ± 10 %). Adult mice (aged between 3 months and 6 months) were divided in three groups: training (n=6 males, n=4 females), test 24h (hours) (n=5 males, n=6 females) and test 7d (days) (n=4 males, n=4 females). All experimental procedures were approved by the Italian Ministry of Health (Authorization n. 512-2018_FC). Alternatives to in vivo techniques were not available, but all experiments were conducted according to principles of the 3Rs.

### BEHAVIORAL TASK

All mice were trained and tested using the behavioral paradigm, step-through inhibitory avoidance (IA). Behavioral procedures were performed in a sound-attenuated room, during the light cycle. Mice were manipulated on the three days previous to training session, and were left in the experimental room from the day before the experiment for habituation. Each mouse was subjected to the task separately. The apparatus consists of an automatic controller and a box (47×18×25 cm, Ugo Basile, Comerio, Italy) which is divided into two separate compartments by a sliding door. The start compartment is brightly-illuminated while the escape one is dark and connected to the shocker. During the training session, every mouse was gently placed in the brightly-illuminated compartment, facing the door and allowing free access to the dark compartment. The apparatus is designed to exploit the natural behavior of mice to move into the dark. For this reason, they rapidly stepped through the door and immediately receive a 0.3 mA mild foot-shock lasting 2 s. Latency was measured using an automated tilting-floor detection mechanism. After the foot-shock, mice were immediately removed from the dark box, and received the 4-OHT injection. The retention test was carried out 24 h or 7 d after the training session. All mice were trained, randomly assigned to be tested 24h or 7d after training. During the test sessions, trained animals were placed again into the light compartment and latency (s) to entered the dark compartment was recorded. The procedure is thus identical to the training session, but mice do not receive shocks. Five minutes is the maximal latency time given for stepping in the dark. At the end of 300 s, mice that do not step through are removed from the apparatus. After removal from the system, all animals received the 4-OHT injection. The latency to step through was considered a direct measurement of memory.

### DELIVERY OF 4-OHT

Mice were handled and got used to needle pain with saline solution daily for at least 3 days prior to the 4-OHT injection. 4-OHT (Sigma H6278) was first dissolved in absolute ethanol to give a final concentration of 20 mg/mL. This stock was then mixed with corn oil (Sigma C8267) at 37 °C in order to get an emulsion. The injectable oil formulation was obtained using the Eppendorf ThermoMixer® C. The emulsion was heated and shaken for 2 hours until the ethanol was entirely evaporated. At this time, the drug was totally dissolved in corn oil and kept at 37°C. 4-OHT (50 mg/kg) was delivered by intraperitoneal (i.p.) injection with a 22 gauge needle.

### EX-VIVO BRAIN PROCESSING

One week after 4-OHT injection, animals were deeply anesthetized with isoflurane (1.5%–2%) and transcardially perfused with 50 mL of ice-cold 0.01 M phosphate buffered saline (PBS) solution (pH 7.6), followed by 75 mL of freshly prepared paraformaldehyde (PFA) 4% (wt/vol, pH 7.6). Brains were extracted and prepared according to the CLARITY/TDE protocol (Chung et al., 2013; Costantini et al., 2015). Immediately after perfusion, brains were post-fixed in PFA overnight at 4°C. The day after, samples were incubated in a hydrogel solution (containing 10% acrylamide (vol/vol), 2.5% bis-acrylamide (vol/vol) and 0.25% VA044 (wt/vol) in PBS) at 4°C for 3 days, allowing a sufficient diffusion of the solution into the tissue. Samples were then degassed, replacing oxygen inside the vials with nitrogen, and incubated in a water bath at 37°C for 3 hours in order to initiate polymerization of the hydrogel. After 3 hours, embedded brains were placed in a clearing solution (containing 4.4 % (wt/vol) sodium dodecyl sulfate (SDS) and 1.2 % (wt/vol) boric acid in ultra-pure water, pH 8.5) at 37°C. Clearing solution was changed every 2-3 days. Specimens were gently shaken throughout the whole clearing period, which typically takes 3-4 weeks. When the samples appeared sufficiently transparent, they were incubated 1 day in PBS with 0.1 Triton-X (PBST, pH 7.6) and 1 day in PBS (pH 7.6), removing the excess SDS. Finally, murine brains optically cleared with serial immersions of mixtures containing 20% and 40% 2-2’ Thiodiethanol (TDE) in PBS, each for 1 day while rotating. The last mixture (40% TDE) was used as an index-matching solution for our imaging (Di Giovanna et al., 2019).

### LIGHT-SHEET MICROSCOPY

The custom-made light-sheet microscope, exploiting RAPID autofocusing, has been described in detail in our previous works (Müllenbroich et al. 2015; Silvestri et al. 2021). In brief, the sample is illuminated from the side using a virtual light sheet created with a galvanometer scanner (6220H, Cambridge Technology), coupled via a 4f system to an air objective (Plan Fluor EPI 10X NA 0.3, Nikon) covered with a protective coverslip. Light emitted from the specimen is detected orthogonally to the illumination plane using an immersion objective corrected for clearing solutions (XLPLN10XSVMP 10X NA 0.6, Olympus). Then, it is bandpass-filtered to isolate fluorescence light and projected by a tube lens onto the chip of a scientific complementary metal-oxide-semiconductor (sCMOS) camera (Orca Flash 2.0, Hamamatsu) operating in rolling-shutter mode to guarantee confocal line detection. During imaging, the sample was fixed in a refractive index-matched quartz cuvette (3/Q/15/TW, Starna Scientific) and moved using a set of high-accuracy linear translators (M-122.2DD, Physik Instrumente). Sample-induced defocus is measured in real-time using RAPID (Silvestri et al. 2021), and correction was implemented by moving the objective with an additional linear translation stage (M-122.2DD, Physik Instrumente). Since an entire mouse brain is too thick to be imaged with our objective, which has a working distance of 8 mm, we imaged the two halves separately, with an overlap thickness of about 1 mm (**Fig. S6**). The entire system was controlled by custom software written in C++, available at https://github.com/lens-biophotonics/SPIMlab.

### IMAGE ANALYSIS

Tiled images acquired with LSM were stitched together using ZetaStitcher (https://github.com/lens-biophotonics/ZetaStitcher). As well as generating a low-resolution view of the entire imaging volume (with voxel side 25 µm), this software includes an application programming interface (VirtualFusedVolume) to access the high-resolution volume. Images were then visualized using FIJI/ImageJ (https://fiji.sc). The two different brain halves were spatially registered with an affine transformation using Advanced Normalization Tools (ANTs) on the low-resolution reconstructions (**Fig. S7**).

### WHOLE-BRAIN CELL DETECTION

Fluorescently labeled neurons were localized in the whole-brain images using BrainCell Finder (Frasconi et al. 2014) (https://github.com/lens-biophotonics/BCFind). In brief, patches of the original dataset (accessed via VirtualFusedVolume) were fed into a UNet with four contraction layers of 3D convolutions with an exponentially increasing number of filters, and four expansion layers of transposed 3D convolutions with a decreasing number of filters. UNet training was carried out with binary cross-entropy loss and Adam optimizer. The goal of this network is to perform semantic deconvolution, that is, to transform the original image into an ideal one in which cell bodies are clearly visible, while other structures such as dendrites and axons are removed. The network was previously trained on a ground-truth dataset in which a human expert has localized the centers of neuronal somata. The training dataset was composed of 221 image stacks for a total volume of approximately 6.8 mm^3^ and 19’166 manually labeled cells. The stacks were randomly selected from different samples and different areas of the brain, to train the network to recognize the large variability in cell shape that can be found across the sample. The images deconvolved by the network are then processed with a standard blob detection algorithm (difference of Gaussians, DoG) to identify the center of bright structures, which in this case are the neurons. The overall performance of the method is evaluated by comparing the list of neuron centers found by the software with the human-annotated ground-truth test set of 61 image stacks for a total volume of approximately 1.9 mm^3^ and 3’383 manually labeled cells. Again, these stacks were randomly selected from different samples and different areas of the brain, to test network performance in different contexts. If two neuron centers from the two annotations (automatic and manual) are closer than 10 μm (approximately half of the average diameter of a neuron), they are considered to be the same cell, that is, a true positive (TP). If a center is present only in the manual annotation, it is considered a false negative (FN), whereas if it is present only in the results of the algorithm, it is considered a false positive (FP). The counting of true positives, false positives and false negatives was carried out using the maximum bipartite matching algorithm (Galil 1986). We evaluated localization performance using the formulas precision = TP/(TP + FP) and recall = TP/(TP + FN), and the F1-score, which is defined as the harmonic mean of precision and recall. For our test set, the precision was 0.84, the recall was 0.74 and the F1-score was 0.78. All annotations of cell positions were performed using Vaa3D (https://github.com/Vaa3D).

### SPATIAL REGISTRATION TO REFERENCE ATLAS

The downsampled version of the whole-brain dataset was spatially registered to the Allen reference atlas using ANTs (Avants et al. 2011), with a sequence of affine and diffeomorphic (that is, symmetric normalization) transformations (**Fig. S8**). In detail, images first underwent a strong gamma correction (with exponent 0.3) to reduce dynamic range and increase the relative contribution of tissue autofluorescence over labeled cells. Then, affine registration between single brains and reference atlas was computed. To reliably assess non-linear deformations introduced by clearing, 3 gross brain regions (cerebellum, hippocampi and olfactory bulbs) were manually segmented. A diffeomorphic transformation was then computed to match the segmented areas with the corresponding ones in the reference atlas. Since this step is needed only to correct large-scale deformations, it is computed on further downsampled versions of the data (100 µm voxel side). Eventually, fine registration to the atlas was computed using a diffeomorphic transformation between the real image and the reference image. This sequence of transformations (including alignment of back and front brain halves) were applied to the point clouds produced by the BrainCell Finder, to represent the position of activated neurons in the reference space. Each cell was then assigned to a selected brain area based on its position. To evaluate registration accuracy, 10 reference points were manually marked in 10 different aligned brains and in the reference atlas (**Fig. S9**). Then, the Euclidean distance between pairs of corresponding points (samples vs atlas) was calculated for each point and each sample (100 measures in total). The root-mean-square average value of the distance (0.30 +/- 0.08 mm, mean +/- s.d.) was considered as the alignment error.

### PARTIAL LEAST SQUARE ANALYSIS

The effect of each experimental group on regional cell counts were evaluated using Task PLS (Krishnan et al. 2011; McIntosh et al. 1996). Briefly, a dummy matrix with as many columns as the number of subjects and as many lines as the experimental classes was prepared, with 1 in the matrix elements corresponding to the right matching subject-class and 0 elsewhere. This was multiplied by the regional count matrix (as many lines as the subjects, as many columns as the brain areas analyzed) normalized by the volume of each area and by the total number of labeled cells per animal, and the product was processed using singular value decomposition. The resulting left and right matrices contained the contrasts and the saliences, respectively. These identify latent variables in the spaces of experimental classes and in the space of cell counts that best explain the variability observed in the data. Statistical error was calculated using the bootstrap technique by resampling 1000 times with replacement. The reported saliences are the raw ones divided by the standard deviation of the bootstrap results, and can thus be interpreted as z-values. In this sense, the dashed lines in Fig.2 represent different significance levels (p < 0.1, p < 0.05, p < 0.01).

### FUNCTIONAL NETWORK GENERATION

The inter-regional correlation (Wheeler et al. 2013) was calculated for each group of mice in order to quantify the co-variation of activated cell counts in 45 brain regions across mice of each group. As fro the PLS, counts normalized by area volume and by total number of labeled cells per animal were used. The functional connections between different regions were identified by imposing a suitable threshold for the inter-regional correlation. The threshold was properly defined to identify cross-correlations with a p-value of less than 0.05. Starting from the correlation matrix and the given threshold value, the corresponding adjacency matrix was computed for each group of mice. Each adjacency matrix defines a corresponding network where the nodes are the 45 brain regions and the links are the functional connections between the regions: positive links correspond to excitatory functional connections and negative links to inhibitory functional connections. The degree centrality of the nodes was calculated by counting the number of connections of each node without considering the sign of the connections, and was normalized on the maximum number of potential connections (in this case 44). The betweenness centrality and the small world coefficient σ were calculated by means of the algorithms betweenness_centrality and sigma of the Python package networkx based on (Brandes 2001; 2008; Freeman 1977) and (Humphries and Gurney 2008), respectively.

---

**SUPPLEMENTARY Table S1.** List of Brain Region.

| ABBREVIATION | LONG NAME | BRAIN REGION |
| --- | --- | --- |
| FRP | Frontal pole | Isocortex |
| MO | Somatomotor areas | Isocortex |
| SS | Somatosensory areas | Isocortex |
| GU | Gustatory areas | Isocortex |
| VISC | Visceral areas | Isocortex |
| AUD | Auditory areas | Isocortex |
| VIS | Visual area | Isocortex |
| ACA | Anterior cingulate area | Isocortex |
| PL | Prelimbic area | Isocortex |
| ILA | Infralimbic area | Isocortex |
| ORB | Orbital area | Isocortex |
| AI | Agranular insular area | Isocortex |
| RSP | Retrosplenial area | Isocortex |
| PTLp | Posterior parietal association areas | Isocortex |
| Tea | Temporal association areas | Isocortex |
| PERI | Perirhinal area | Isocortex |
| ECT | Ectorhinal area | Isocortex |
| OLF | Olfactory areas | Olfactory areas |
| HIP | Hippocampal region | Hippocampal formation |
| RHP | Retrohippocampal region | Hippocampal formation |
| CLA | Clastrum | Cortical subplate |
| EP | Endopiriform nucleus | Cortical subplate |
| LA | Lateral amygdalar nucleus | Cortical subplate |
| BLA | Basolateral amygdalar nucleus | Cortical subplate |
| BMA | Basomedial amygdalar nucleus | Cortical subplate |
| PA | Posterior amygdalar nucleus | Cortical subplate |
| STRd | Striatum dorsal region | Striatum |
| STRv | Striatum ventral region | Striatum |
| LSX | Lateral septal complex | Striatum |
| sAMY | Striatum-like amygdalar nuclei | Striatum |
| PALd | Dorsal pallidum | Pallidum |
| PALv | Ventral pallidum | Pallidum |
| PALm | Medial pallidum | Pallidum |
| PALc | Caudal pallidum | Pallidum |
| DORsm | Thalamus, sensory-motor | Thalamus |
| DORpm | Thalamus, polymodal association | Thalamus |
| PVZ | Periventricular zone | Hypothalamus |
| PVR | Periventricular region | Hypothalamus |
| MEZ | Hypothalamic medial zone | Hypothalamus |
| LZ | Hypothalamic lateral zone | Hypothalamus |
| ME | Median eminence | Hypothalamus |
| MB | Midbrain | Midbrain |
| P | Pons | Pons |
| MY | Medulla | Medulla |
| CB | Cerebellum | Cerebellum |

**Table S2.**
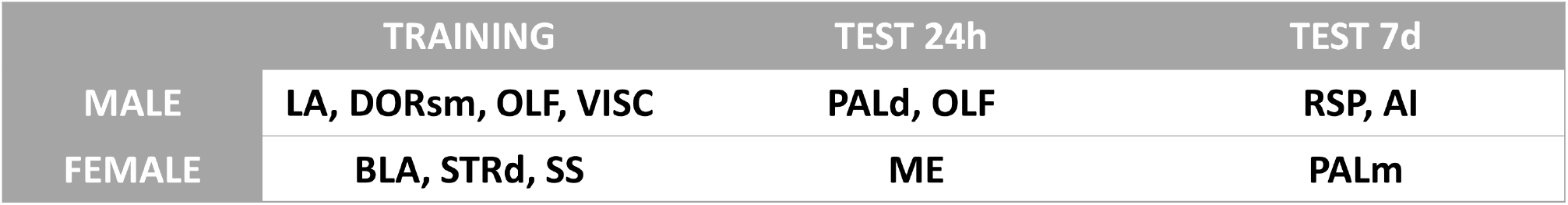
Putative functional connectivity hubs in male and female mice.

|  | TRAINING | TEST 24h | TEST 7d |
| --- | --- | --- | --- |
| MALE | LA, DORsm, OLF, VISC | PALd, OLF | RSP, AI |
| FEMALE | BLA, STRd, SS | ME | PALm |

## SUPPLEMENTARY MATERIAL

**Figure S1:**
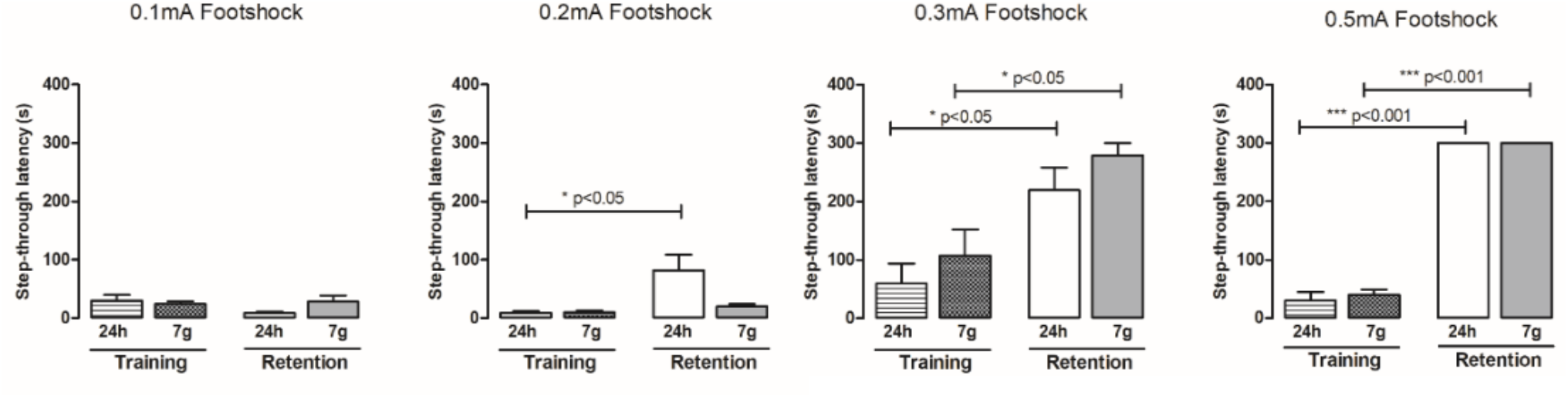
Evaluation of four shock intensities for the optimization of step-through paradigm. Comparisons of acquisition and retention latency times are analyzed by 1-way ANOVA, followed by Bonferroni’s post-hoc comparisons tests. Data are expressed as means ± SEM of 7 to 10 animals for each group.

**Figure S2:**
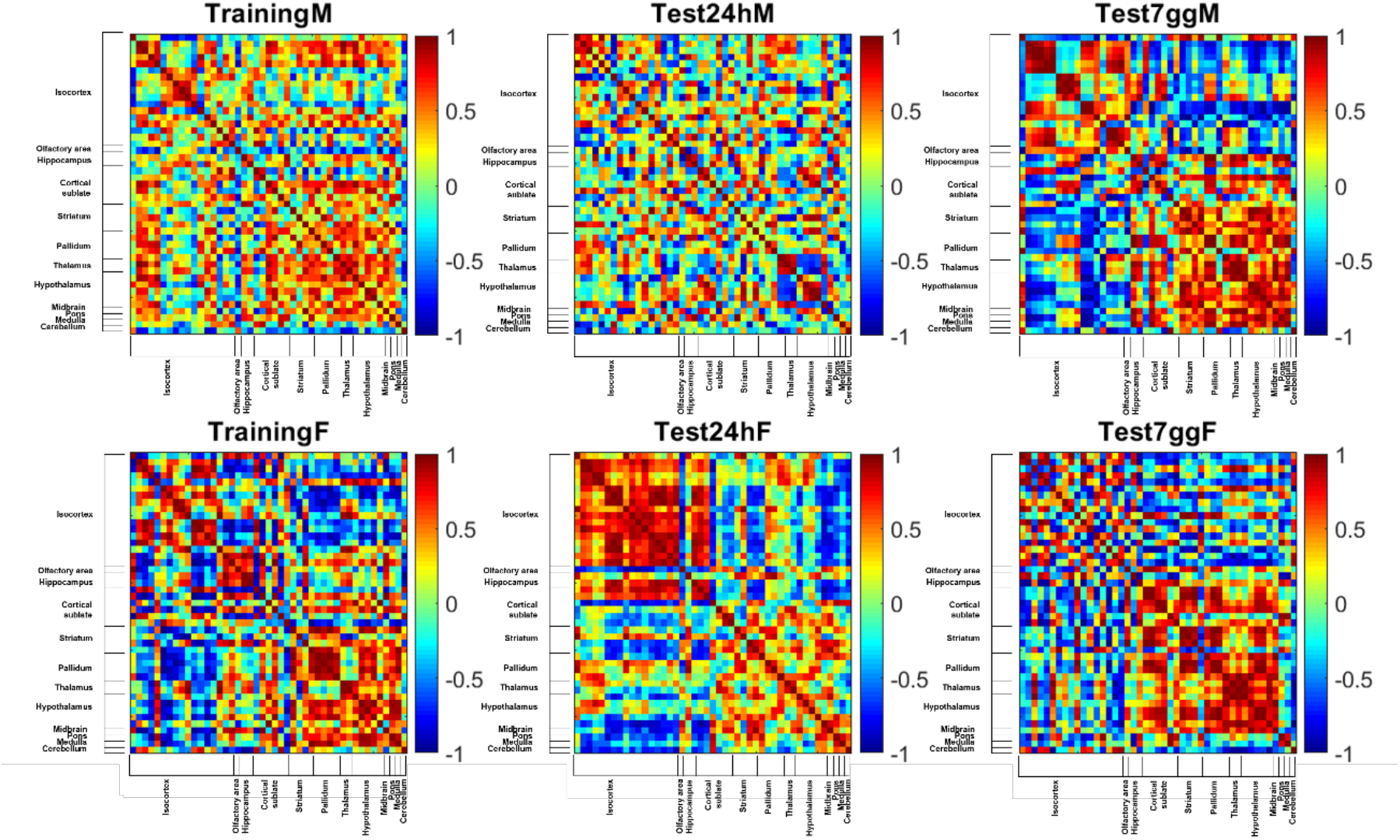
Generation of Pearson correlation heatmap following the three experimental time points. Matrices showing Pearson correlations between cFos expression of different brain areas at training, test 24h, and test 7d in both sexes. Axes correspond to 45 brain regions shown in Tab S1. Color bar on the left reflects the value of Pearson coefficient.

**Figure S3:**
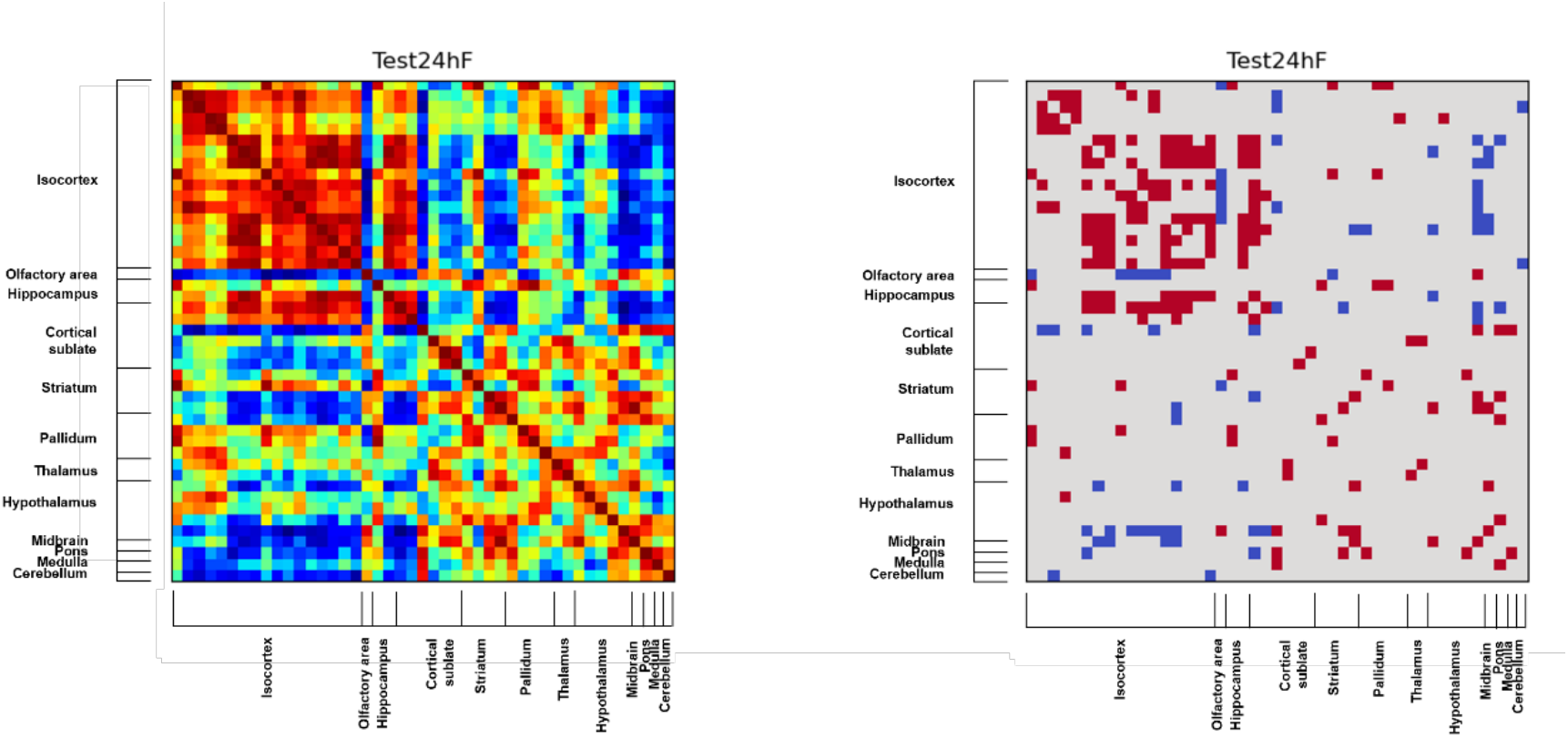
Generation of Matrix connectivity. Matrix on the left represents a Pearson correlation heatmap while matrix on the right is the same in which only correlation with a p value< 0.05 are considered. Axes correspond to 45 brain regions shown in Tab S1.

**Figure S4:**
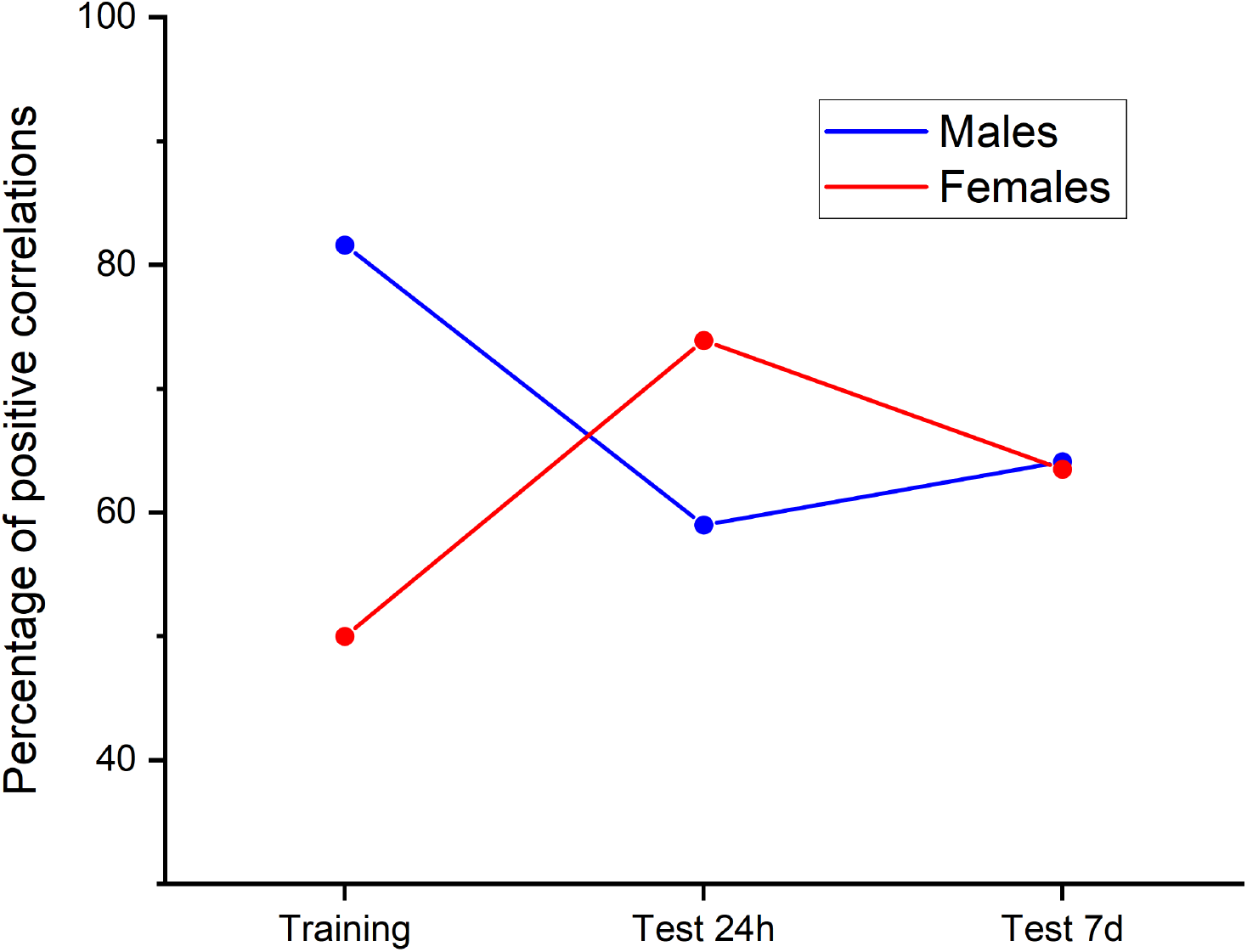
Fraction of positive correlations. Graph that shows the percentage of positive correlations in the functional connectome for each experimental class and for both sexes.

**Figure S5:**
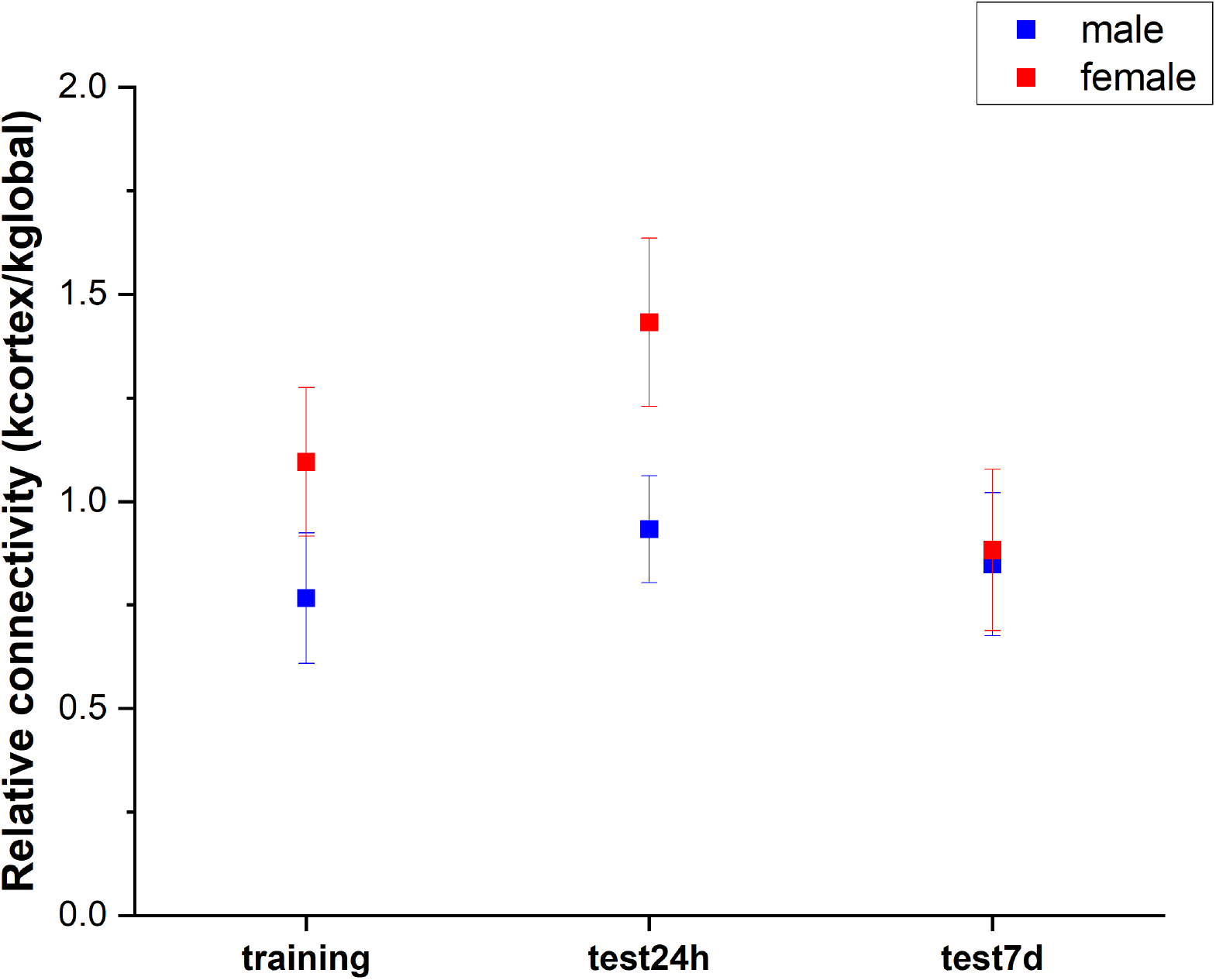
Cortex connectivity. Relative connectivity is calculated by the ratio of the degree mean value of the isocortex and degree mean value of the entire network. For each experimental class, relative connectivity is shown for both sexes.

**Figure S6:**
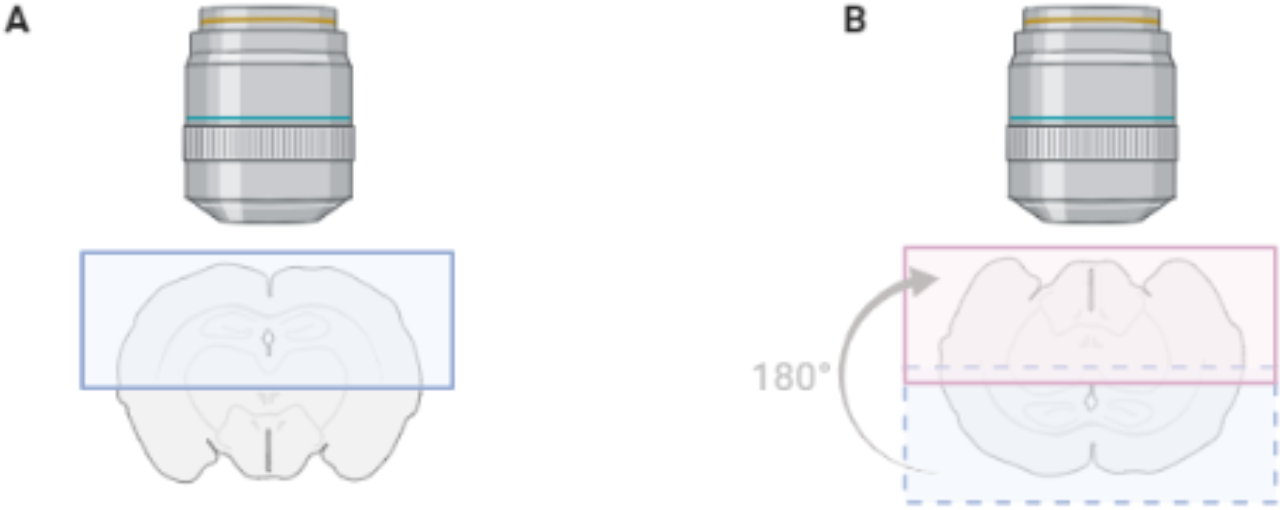
Schematic representation of two-halves imaging. The brain half, closer to the objective (in blue), was imaged first (A). Then, the sample was rotated and the other half (in red) was imaged (B).

**Figure S7:**
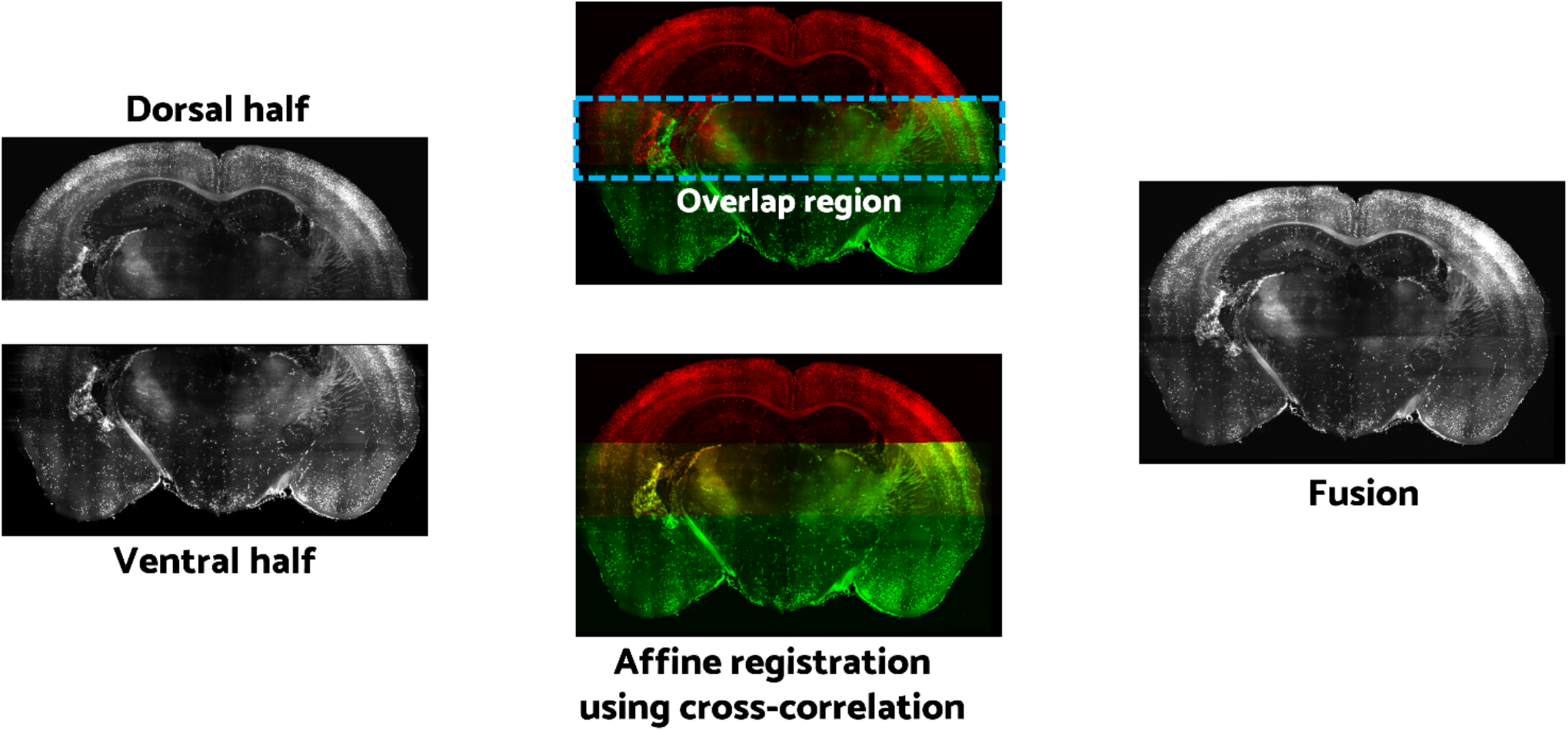
Schematic representation of the fusion of two-halves brain. After the imaging part, dorsal and ventral brain halves are overlapped and fused using an affine transformation. In the middle, brain half colored in red corresponds to the dorsal half. Green one corresponds to the ventral half. The overlap region is shown in yellow.

**Figure S8:**
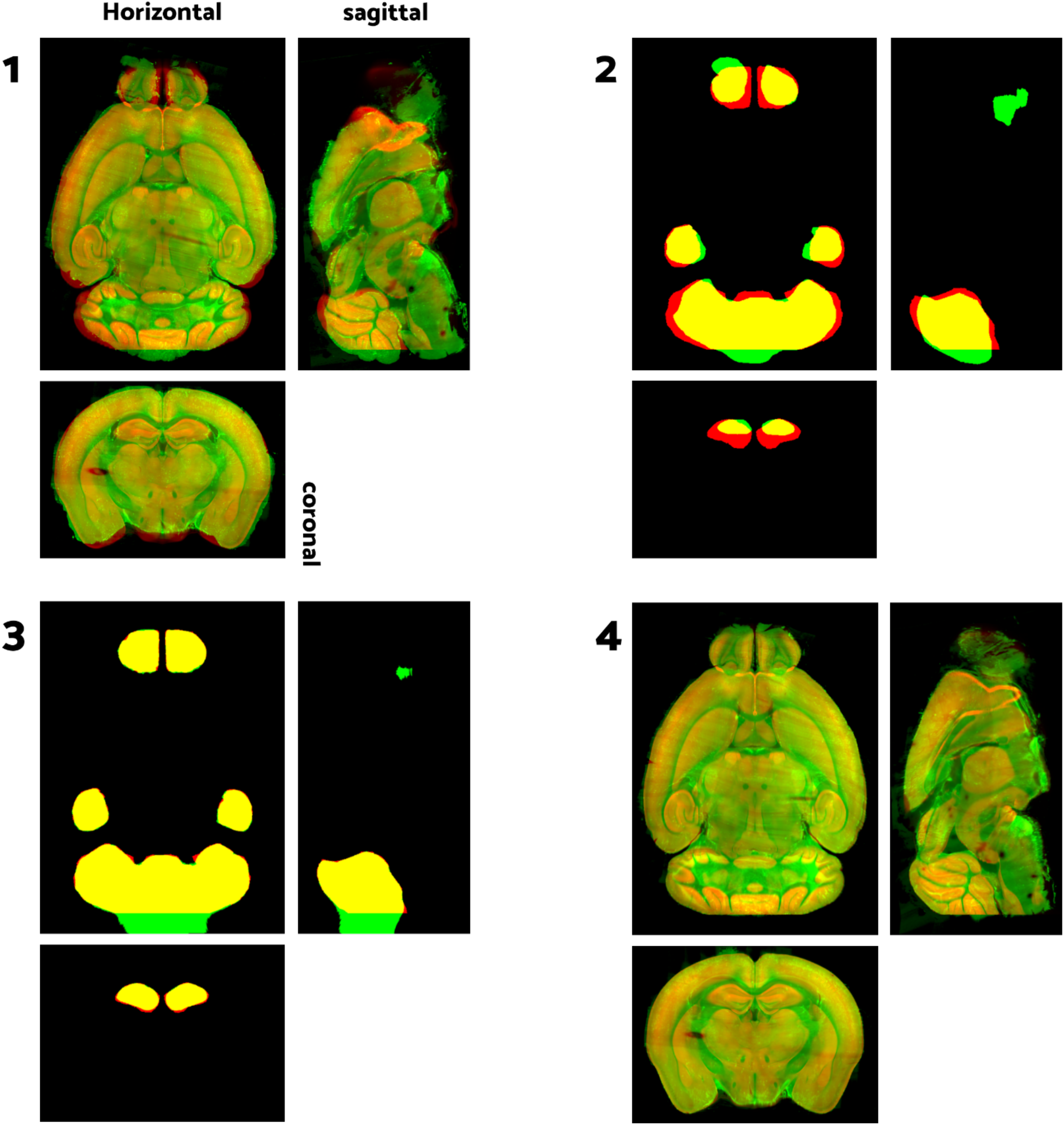
Atlas registration. Four steps are necessary for registering whole-brain fluorescence datasets with standard reference. Reference Atlas and relative masks are in red, while LSM images and masks are in green. (1) The first step is a gross registration of LSM datasets with images of Reference Atlas through an affine transformation. (2) From these images, binary masks of 3 main anatomical regions (olfactory bulbs, hippocampi, and cerebellum) are extracted. (3) Large distortions are corrected by running a non-linear registration on these masks. (4) Using alignments computed from the masks, a final non-linear registration between the sample and the atlas is found.

**Figure S9:**
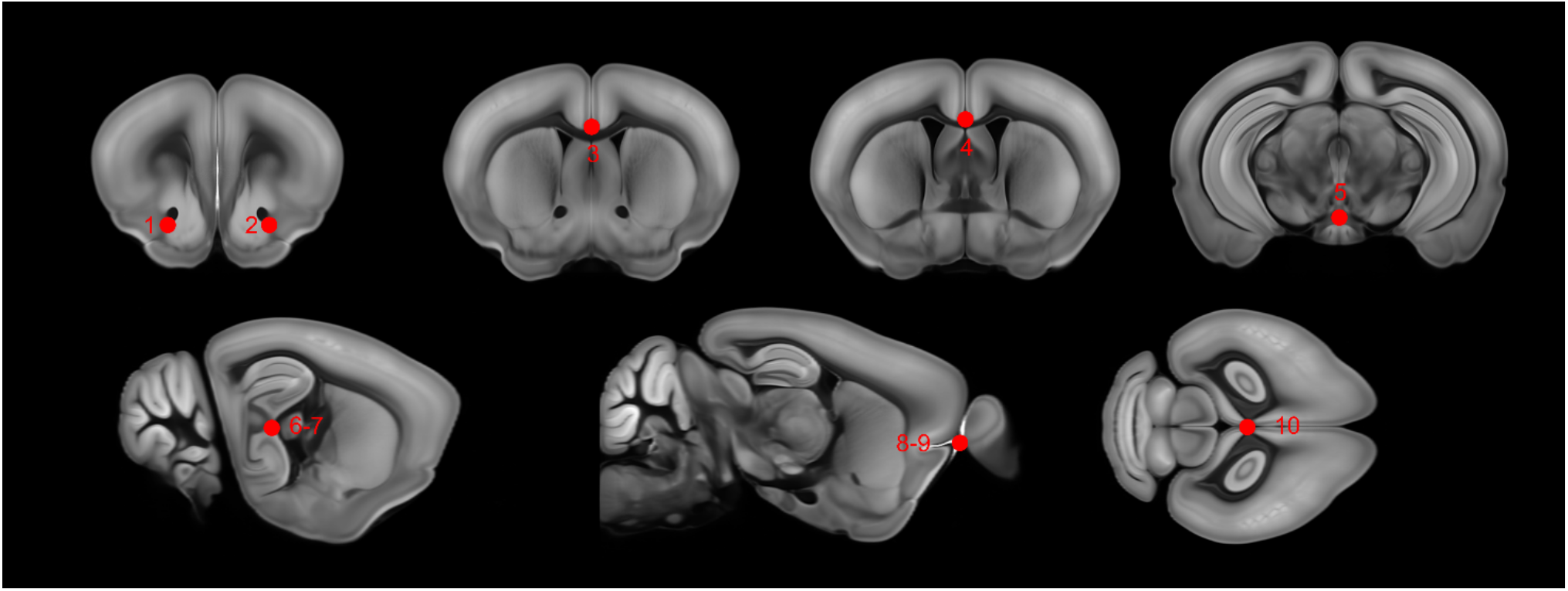
Evaluation of Atlas registration accuracy. In the picture, six slices of the Reference Atlas (in the coronal, sagittal, and horizontal plane) are shown. The image reveals the annotations of ten red points necessary to calculate the registration accuracy. In the fifth and sixth slices, red points are annotated in each hemisphere.

## SUPPLEMENTARY VIDEO

***Video V1:* Whole-brain video**. Navigation in the 3D rendering of a whole TRAP mouse brain. The video displays the entire brain at low resolution, to be visualized on the screen. The whole brain is then shown in sequential coronal and sagittal planes. Finally, a representative zoom-in is shown at the native resolution of the light sheet microscope. Cell morphology can be recognized demonstrating the sensibility of the system.

***Video V2:*** Coronal stacks of the 24 mouse brains analyzed in this study. White spots represent all nurons activated in the different experimental classes. Train is the abbreviation for training group, Test 24 and Test 7 is test after 24 hours and 7 days after training. M and F after the name indicate male and female.

***Video V3:*** Z-stack showing the images from a TRAP mouse brain acquired with light sheet microscopy (left) and the corresponding heatmaps displaying cell density (right).

## Author contributions

L.S. conceived research and coordinated the project; A.F. performed animal experiments, prepared and imaged samples; M.B.P. designed behavioral experiments; G.M. and C.C. developed BRANT-related software; F.S.P. contributed advanced imaging tools; I.C. contributed advanced clearing methods; L.C. and D.F. performed network analysis; B.A.S., A.F. and L.S. analyzed data; A.F. and L.S. wrote the manuscript with contribution from all other authors.

## Acknowledgments

The Authors are thankful to Antonino Paolo di Giovanna, Barbara Rani and Alessia Costa, University of Florence, for lab support in setting up protocols with tissues and animals.

